# Distinct cortical encoding of acoustic and electrical cochlear stimulation

**DOI:** 10.1101/2025.08.01.668170

**Authors:** Ariel Edward Hight, Michele N. Insanally, Julia K. Scarpa, Yuko Tamaoki, Rohit Makol, Yew-Song Cheng, Michael Trumpis, Jonathan Viventi, Mario A. Svirsky, Robert C. Froemke

## Abstract

Cochlear implants are neuroprosthetic devices that restore hearing and speech comprehension to profoundly deaf humans and represent an exemplar application of biomedical engineering and research to clinical conditions. However, the utility of these devices in many subjects is limited, largely due to lack of information about how neural circuits respond to implant stimulation. Recently we showed that deafened rats can use cochlear implants to recognize sounds, and that this training refined the responses of single neurons in the primary auditory cortex. Here we asked how local populations of cortical neurons represent acute implant stimuli, using electrode arrays we developed for cortical surface recordings for micro-electrocorticography (µECoG), a form of intracranial electroencephalography (iEEG). We found that there was a coarse, non-random spatial organization with limited evidence for consistent, sharply graded cochleotopy across recording sites, relative to a clearer tonotopic spatial representation in normal-hearing rats. Single-trial iEEG responses to acoustic inputs were more reliable than responses to cochlear implant stimulation, although stimulus identity could be successfully decoded in both cases. However, the spatio-temporal response profiles to acoustic vs cochlear implant stimulation were substantially different. Decoders trained on acoustic responses showed essentially zero information transfer when tested on electrical stimulation responses in the same animals after deafening and cochlear implant stimulation. Thus, while acute cochlear implant stimulation evoked spatially non-random cortical activity with coarse cochleotopic structure, the dynamics of network activity were substantially different from those evoked by acoustic stimulation, with possible implications for perceptual similarity that remain to be tested.

## Introduction

Cochlear implants are neuroprosthetic devices that restore hearing and provide users with an ability to attain open-set speech perception without other aids such as lip-reading (Djourno and Eyries, 1957; Simmons et al., 1965; Michelson, 1971; House and Urban, 1973; Eddington et al., 1978; Hochmair and Hochmair-Desoyer, 1981; Wilson et al., 1991; Clark, 2006; Svirsky, 2017). Cochlear implants function by inserting a single and flexible shank of electrodes along the cochlear spiral (generally 12-22 channels in human subjects), to stimulate primary afferent neurons of the auditory system and bypass pathologies of deafness. Individual electrodes deliver trains of electrical pulses, providing spectral cues as a function of location along the topographical axis of the cochlear spiral, mimicking the encoding of sound in a functional and healthy cochlea. Over 1 million users have been implanted world-wide, making it the most widely adopted type of brain-computer interface, surpassing vagus nerve stimulators, deep-brain stimulation electrodes, and artificial retinas (Hays et al., 2013; Dawson et al., 2016; Ayton et al., 2020; Zeng, 2022; Johnson et al., 2024). Cochlear implants are thus the gold-standard for neuroprosthetic device use in terms of performance, safety, and ability to fine-tune or personalize the programming of each device to individual users.

Despite their wide adoption in human subjects over the last several decades, there is limited understanding of how these devices restore functionality and the sense of hearing to users. One long-standing question has been how the central auditory system responds to and interprets signals resulting from electrical stimulation of auditory nerve fibers. It is unknown to what degree emulating the patterns evoked by acoustic stimulation in normal hearing users is important for the auditory system to interpret patterns of evoked neural activity and inform downstream auditory perception. Many structures of the central auditory system are organized tonotopically, following the frequency alignment of the cochlear basilar membrane (Walzl and Woolsey, 1946; Merzenich et al., 1973; Polley et al., 2007). This topographic mapping might facilitate the encoding of auditory stimuli. Previous studies have demonstrated cochleotopic organization in auditory cortex of cochlear implant animals (Bierer and Middlebrooks, 2002; Johnson et al., 2016). Here, our aim was not to establish spatial organization per se, but to test whether acoustic and electrical stimulation produce overlapping or transferable cortical population representations under a common analysis framework. Previous psychophysical studies in human implant users have provided indirect evidence for perceptual differences (McDermott, 2004) and similarities (Vermeire et al., 2013), but direct neural evidence comparing representations in the same subjects has been lacking. This question has profound implications for how we design and program cochlear implants, or more generally other brain-computer interface devices.

Studies in non-human animals are required to understand the neural basis of cochlear implant use and the relation to spatiotemporal representations in the auditory system. Responses to cochlear implant stimulation have been studied in a number of species generally in non-behaving animals under anesthesia (Klinke et al., 1999; Bierer and Middlebrooks, 2002; Middlebrooks and Bierer, 2002; Fallon et al., 2014; Johnson et al., 2016, Adenis et al. 2024), but occasionally in awake or behaving animals (Vollmer and Beitel, 2011; Keating et al., 2013; Johnson et al., 2016). Over the last decade we have examined cochlear implant responses in deafened rats (King et al., 2016; Glennon et al., 2019), where we showed that animals can be trained to report the activation of specific implant channels. Behavioral training with the cochlear implant led to substantial plasticity within the deafened rat auditory cortex, modifying synaptic receptive fields and adjusting excitatory-inhibitory balance at the level of single neurons. Prior to training, the responses of single cells to implant stimulation were erratic and inhibition was mismatched relative to excitation. After training, inhibition became aligned with excitation across the array of implant electrode channels (Glennon et al., 2023). However, intracellular recording in vivo is infeasible in human subjects. Thus other approaches more amenable to clinical use are required, to measure neural coding of cochlear implant signals and enable translation of results from non-human studies to improve implant performance by human subjects.

Here we aimed to determine the spatiotemporal representations of implant channels beyond the responses of single neurons, but instead across a broader extent of rat auditory cortex. We take advantage of improvements in micro-electrocorticography (µECoG), a sub-type of intracranial electroencephalography (iEEG) that generally uses grid electrodes placed epi- or sub-durally over the cortical surface to measure local electrical activity (Insanally et al., 2016; Woods et al. 2018). iEEG recordings have high temporal precision, greatly improved signal-noise ratios, and better spatial resolution than conventional scalp EEG recordings (Abdi-Sargezeh et al., 2023; Janiukstyte et al., 2023). Signals reflect local electrical fields generated by aggregate neural activity in the regions adjacent to recording sites (Jasper and Penfield, 1949; Chang, 2015; Fukushima et al., 2015). iEEG has been used for studies of human brain organization particularly of speech and language centers, but these recordings are necessarily done in epileptic subjects to help locate seizure foci before surgical resection (Nourski et al., 2013; Mesgarani et al., 2014; Tang et al., 2017; Beynon et al., 2021; Norman-Haignere et al., 2022). Non-human animal studies remain essential for determining how sensory inputs are represented and processed in subjects without overt neurological conditions. To this end, we have designed and manufactured a novel and flexible 60-channel cortical surface iEEG array for adult rats and validated that we can measure auditory responses and cortical map topography in normal hearing animals with these arrays (Insanally et al., 2016; Trumpis et al., 2017; Woods et al. 2018). Using these custom-fabricated iEEG grid electrodes, we now ask how auditory cortex responds to implant stimulation in untrained or trained deafened adult rats, and how different evoked signals and/or mesoscale tonotopic organization is in cochlear implant users vs normal-hearing animals.

## Materials and Methods

### Surgical procedure for iEEG recordings

All animal procedures were performed in accordance with National Institutes of Health standards and were conducted under a protocol approved by the NYU Grossman School of Medicine Institutional Animal Care and Use Committee. We used a custom array consisting of 61 passive electrodes spaced 406 μm apart in an 8 x 8 grid, with three corner electrodes omitted and the fourth corner electrode intentionally unexposed to test encapsulation, reducing the number of active recorded sites to 60 (Insanally et al., 2016). iEEG grids were fabricated using a standard flex-PCB processing technique by Microconnex Flex Circuits (Snoqualmie, WA). All procedures and experiments were carried out in a sound-attenuation chamber and under anesthesia.

A specific surgical protocol was developed for implantation of cortical surface iEEG in rats. Sprague-Dawley rats 4–6 months old were anesthetized with ketamine (40 mg kg−1, intramuscular injection) and dexmedetomidine (0.125 mg kg−1, intramuscular injection), or pentobarbital (50 mg kg−1, intraperitoneal injection). The head was secured in a custom head-holder that left the ears unobstructed. A longitudinal incision was made along the midline to expose the skull. Five bone screws were inserted into the skull around the point of entry of the electrode array to help anchor the dental cement (C & B Metabond Quick! Luting Cement). After reflecting the right temporalis muscle, a 5 mm × 5 mm craniotomy was made on the right temporal skull to expose the brain, and a sterilized iEEG array was epidurally placed over the left hemisphere core auditory cortex, located using vasculature landmarks. A thin silver wire was soldered between the headstage and the skull screws to be used as ground and reference.

### Surgical procedure unilateral cochlear implantation of the right ear

Cochlear implantation procedures are similar to our past studies (King et al., 2016; Glennon et al., 2023). 8-channel animal cochlear implant arrays (HL08) were provided by Cochlear Americas. The array contained platinum-iridium band electrodes coated in silastic and connected to a nine-pin Nanonics connector (TE Connectivity, Berwyn, PA) with a single, additional extracochlear ball ground. The ipsilateral pinna was pulled forward and secured with a hemostat, the head rotated laterally, and the post-auricular junction between the ear canal and the sternocleidomastoid muscle (SCM) is identified as the initial incision site. An incision was made and the superficial fascia of the neck was dissected to identify the facial nerve i.e., cranial nerve (CN) VII. Minor bleeding was controlled using hemostatic epinephrine-soaked cotton pellets (Epidri pellets; Pascal International, Bellevue, WA), applied with light pressure. The SCM and posterior belly of the digastric muscle (PBD) were dissected from the tympanic bulla (TB) ventral and rostral to the trunk of CN VII. The TB was cleared of muscle and periosteum; the periosteum of the bulla was kept in normal saline and used later to seal the cochleostomy site. The drilling of the TB was begun ventrorostral to the trunk of CN VII with a 0.5 mm diamond burr and continued dorsally with care taken to avoid injuring CN VII, until the SA overlying the RW was fully visualized. Any remaining tissue or debris was removed with microforceps before the cochleostomy was performed.

Prior to performing the cochleostomy and inserting the array, the array lead and connector were secured. The post-auricular incision was expanded dorsally toward the skull. An area 4-5 mm in diameter was cleared and cleaned to expose the occipital skull, and the connector was attached perpendicular to the skull using C&B-Metabond (Parkell, Edgewood, NY) and bone screws. The lead to the electrode array was then sutured to the trapezius muscle, allowing enough lead to remain free to facilitate motion required for array insertion. The lead to the separate ground electrode was similarly secured into small muscle pockets in the trapezius. The cochleostomy site was identified ∼0.5 mm directly below the lip of the RW in the basal turn of the cochlea, identified by both the stapedial artery and the cochlear promontory in the tympanic space. The site was gently drilled with a 0.1 mm diamond burr, and the array was inserted into the scala tympani without resistance using AOS forceps (Cochlear, Sydney, Australia) until all of the platinum-iridium contacts were within the scala tympani. We required that all eight electrodes pass through the cochleostomy, confirming that the array was inserted in the direction of the apex. The array occludes most, if not all, of the drill site, but to minimize postsurgical perilymphatic leak strips of periosteum taken from the bulla were placed around the implant to seal the site, followed by application of high-grade cyanoacrylate (Surgi-lock 2oc; Meridian Animal Health, Omaha, NE). The remaining lead was cemented into the bulla with C&B-Metabond (Parkell). Before closure, a small square of gelfoam with dexamethasone was left on the root of the facial nerve to prevent inflammation and heal any minor damage that may have occurred.

### Bilateral sensorineural hearing loss

The deafening procedure (mechanical only, as described and validated in Glennon et al. 2023) was identical to the cochlear implantation procedure, with the exception of the array being removed before closure. Following both array and gelfoam removal, the cochleostomy site was closed with a trapezius muscle or periosteum graft, followed by 2-octyl cyanoacrylate (i.e., the array is removed from the cochlea). Both ears were deafened in this manner, but a functional array remained in the right ear for acute electrical stimulation. We did not obtain ABR measurements, hair-cell counts, or behavioral confirmation of deafening in the animals used for the present iEEG dataset; thus, support for deafening efficacy in this cohort relies on prior validation of the same procedure in separate cohorts (Glennon et al., 2023).

### Behavioral training for tone and implant channel detection

Four animals with iEEG recording electrodes were behaviorally trained to detect target tones. Rats were food restricted and trained on a self-initiated, auditory go/no-go task (Froemke et al., 2013; Martins and Froemke, 2015; King et al., 2016). Animals nosepoked in a designated port to initiate trials and are trained to nosepoke in a different port if the target tone was presented (22.6 kHz, any intensity) or withhold from nosepoking if a nontarget (foil) tone was presented (0.5-32 kHz excluding 4 kHz if 4 kHz was the target, or 8-45.3 kHz excluding 22.6 kHz if 22.6 kHz was the target, at 0.5-1 octave intervals and at any intensity). A sugar pellet reward was given for correct nosepokes within 2.5 s of target-tone presentation, whereas a 7 s timeout was given if the animal incorrectly nosepoked for foil tones. The four rats that achieved >70% target-tone hit rate and d’ ≥1.7 were included for further testing and implantation. A second subset of three animals was behaviorally trained to detect cochlear implant channel 3 (N=2) or channel 4 (N=1); procedures were the same for tones except that a programmable clinical cochlear implant processor was used. These animals were trained for 3, 9, and 13 days.

### Stimulus presentation for cortical sensory mapping in normal hearing rats

Tone presentation was similar to our previous study of iEEG responses in normal-hearing rats (Insanally et al., 2016). Pure tones 50 ms in duration with 2 ms cosine-squared ramps ranging from 0.5-32 kHz (0.5 octave spacing) were generated using an auditory processor (TDT System III RZ6) and presented at a pseudorandom sequence at a rate of 1.25 Hz to the contralateral (right) ear using a calibrated free-field speaker (MF1 Multi-Field Magnetic Speaker, Tucker-Davis Technologies). Each tone was repeated 30 times at 70 dB SPL. The calibrated speaker exhibited <1% harmonic distortion and a flat output in the frequency range used.

### Stimulus presentation for cortical sensory mapping in cochlear implanted rats

Electrical stimulation was delivered by an off-the-shelf Nucleus Freedom system speech processor (Cochlear) in which its transmitter coil drove a CI24RE implant emulator, where its output was connected to the implanted electrodes. The implant emulator is a standard clinical cochlear implant that is mounted in a plastic box with a DB-25 connector (Cochlear). We created a pigtail wire with a DB-25 connector and an Omnetics/Nanonics connector (Omnetics/TE Connectivity) to couple the emulator to the skull-anchored connector. The skull-anchored connector is also an Omnetics/Nanonics connector that is directly attached to the implanted array. All stimuli were based off standard parameters for cochlear devices including charge-balanced biphasic pulses with 8 µs interphase gaps and 25 µs/phase (total pulse width = 58 µs), presented at 900 pulses per second and configured in monopolar configuration where currents were driven between a single intracochlear electrode and the external ground electrode. For about half of tested animals (N=3), auditory stimulation activated the implant through the microphone of the speech processor via tones presented to each individual electrode’s frequency allocation. The remaining cochlear implanted animals (N=4) were stimulated directly; the CI24RE implant emulator was driven by a Freedom system speech processor connected through the Freedom Programming Pod to a personal computer running the Custom Sound EP software (Cochlear). The Custom Sound EABR function was used (5 stimulation pulses presented at 900 pulses per second) to program and deliver stimuli to the implant.

### Cochlear implant programming

Impedance and threshold measurements were obtained intraoperatively using Custom Sound EP (Cochlear) and were used for the initial programming of the sound processor. Electrically-evoked compound action potential (ECAP) thresholds were obtained and used to set the maximum stimulation level and the minimum stimulation level was set to 30 ‘clinical units’ below the maximum level, equivalent to 4.7 dB in the CI24RE implant emulator (Azadpour and McKay, 2012). All additional settings typically deployed in a clinical setting, such as ADRO, were turned off (Custom Sound, clinical programming software for Nucleus Freedom). In some animals, electrodes in the 8-channel array were non-functional prior to implantation (e.g., high impedance), and therefore those animals contributed fewer than eight usable implant channels to subsequent analyses (animals and electrodes are indicated in Supplemental Fig. 3). In these animals, all analyses removed these non-functional electrodes while preserving the cochleotopic identity of functional electrodes.

### ECAP measurements

In a separate cohort (N=3), we obtained ECAP measurements in deafened and implanted rats that were not otherwise used for iEEG recordings. These animals were acutely implanted with an 8-channel cochlear implant and measured for ECAPs by stimulating and recording from the array of implanted electrodes, with an additional reference electrode implanted subdermally in the neck, using a cochlear implant emulator and a custom software program written in MATLAB and Python and using the Nucleus Implant Communicator (NIC, Cochlear Ltd.). All ECAP measures used the “ABCD” artifact removal technique (Brown et al. 1990) with a probe-masker interval of 0.5 ms, and a similar artifact removal for forward masking measures that was modified from the ABCD technique to compensate for different masker electrode configurations (Azadpour et al. 2021). The sense/recording electrode was selected as two channels apical to the probe; if that channel was not available, then the sense was selected as two channels basalward. In cases where the masking electrode was the same as the sense electrode, the sense was switched to the electrode closer to the probe, and the corresponding masked ECAP was scaled relative amplitudes of the probe-evoked masker-free ECAP measured at the two different sense electrodes. After impedance measurements were made to confirm the device was correctly implanted, configured, and within safe/compliant limits, ECAP thresholds were estimated for each electrode by manually stepping up and down stimulation levels specific to the manufacturer’s programming units by steps of 5, until evoked ECAP amplitudes were of ∼100 µV. Next, a forward masking paradigm was implemented to measure channel interaction and temporal recovery. For channel interaction, probe electrodes were preceded by a masker electrode by 0.5 ms, and masker electrode was roved from channel 1-8 to extract spatial tuning functions for each probe electrode. For temporal recovery, a masker stimulus was delivered on the electrode as the probe and the masker-probe interval was roved to capture temporal tuning functions for each probe electrode.

### Processing of cortical iEEG recordings

All signal processing and analyses were performed using custom MATLAB scripts (MathWorks, MA). To determine whether trials were functional or contained artifacts, raw measurements were first filtered (2-150 Hz, 6^th^ order bandpass IIR filter) and then root mean squared power (rms) was computed for each channel. All trials with rms 20-300 µV were considered functional; power less than 20 µV corresponded with pre-amplifier saturation, while power greater than 300 µV consistently indicated intermittent corruption from non-neural sources. To then process functional recordings, we first used a notch filter to remove 60 Hz noise and 90, 120, 180 Hz harmonics. Recorded signals were then downsampled from 20 to 2 kHz using the decimate function (MATLAB), performing also the function of low-pass (anti-alias) filter and thereby rejecting any artifacts resulting from electrical stimulation delivered by cochlear implant electrodes.

Event-related potentials (ERPs) were extracted from raw downsampled recordings by further bandpass filtering (2-100 Hz) using a digital zero-order 6^th^ order Butterworth filter. Magnitudes of evoked ERP transients were measured by first rectifying measurements using the MATLAB abs() function and then subtracting the maximum of the baseline amplitude in the 50 ms pre-stimulus period (averaged over ±1 timestep) from the maximum of evoked amplitude in the 50 ms post-stimulus period (also averaged across three trials).

High gamma responses were extracted from raw downsampled recordings by further bandpass filtering (70-140 Hz) using a digital zero-phase 6^th^-order Butterworth filter. Next, signals were rectified and smoothed with a 20 ms sliding window. Resulting signals that exceeded 5 times the 90^th^ percentile were interpolated. Evoked high gamma magnitudes were extracted as for ERPs.

### Estimation of best frequency

All analyses of tone-evoked responses were restricted to the range of 1.4-32 kHz tones. Best frequency was estimated by computing the center of mass of tone-evoked responses at 70 dB SPL. The response mass function was defined by the mean vector after projecting all responses on a circular domain where vector angles are represented by tone frequencies, to prevent biasing the center of mass towards the interior of the 1.4-32 kHz stimulus range.

### Principal component analysis (PCA)

Evoked responses were analyzed by including both temporal (i.e., *R* post-stimulus responses) and spatial (i.e., *S* responsive sites) variables. These measurements were concatenated across individual trials, *T*, to create a 2-dimensional *T* (rows) x *R*-*S* (columns) matrices. PCA was then performed and *R*-*S* vectors were compressed using singular value decomposition to obtain a reduced-rank approximation of the original matrix. PCA was also performed on variations of our data including only spatial *P* values (i.e., including only evoked magnitudes) or only temporal *R* values (i.e., averaging responses across all recording sites). The top 15 ranked projections were preserved for further analysis, reducing the datasets into a *T* (rows) x 15 (column) matrix. Fifteen components were retained because this modestly exceeded the number of unique stimuli (10 tones) while avoiding unnecessary dimensionality. Scree plots showed that explained variance entered a near-linear, low-slope regime beyond this point, indicating diminishing returns for additional components.

### Tensor component analysis (TCA)

We applied a canonical polyadic (CP) decomposition to a 3-dimensional model, enabling us to treat spatial and temporal aspects of iEEG recordings across multiple trials as linearly independent (orthogonal) dimensions (Kolda and Bader, 2009; Williams et al., 2018). Naive feature vectors were constructed by arranging all data into a three-dimensional matrix composed of length-*R* post-stimulus responses, *S* responsive sites, and *T* stimulus trials resulting in a *R* x *S* x *T* matrix. CP decomposition was performed (Bader et al., 2023) for 15 latent factors, the number 15 chosen to exceed the number of tone or implant stimuli and to parallel the data reduction performed by PCA. Then the resulting estimate of the *T* factors for each of the 15 components reduced our datasets into *T* (rows) x 15 (column) matrices. For cross-modal analyses, TCA was used because it constrains latent factors into separable spatial, temporal, and trial dimensions. This allowed spatial and temporal factors learned in the normal-hearing condition to be held fixed while re-optimizing only the trial factors in withheld normal-hearing data or cochlear-implant-evoked data, thereby providing an interpretable test of within-modality recovery and cross-modal generalization.

### Linear discriminant analysis (LDA)

We trained a decoder for predicting stimulus identity from single trial recordings by developing a supervised linear classifier. Each decoder was retrained and retested 1,000 times to average decoder performance across randomizations for combinations of trials used for training and test sets. Specifically, each bootstrap replicate was randomly partitioned to include 13 trials per stimulus for each training set and the remaining trials being used to test the decoder. LDA was applied to estimate the stimulus-likelihood map of the PCA- or TCA-compressed feature space. Naive feature vectors from the training set were normalized and compressed according to the learned transformations, and the maximum likelihood stimulus (i.e., the identity of the tone frequency or active cochlear implant electrode) was estimated for each response. Absolute accuracy of the decoder was computed as the average accuracy over all stimuli was estimated by the total proportion of correct predictions. In addition to absolute accuracy, we also computed the mean of the error distances between the predicted and actual stimulus, i.e., octaves or number of electrodes.

### Linear mixed effects modeling

Group comparisons between normal-hearing and cochlear implant conditions were assessed using linear mixed-effects (LME) models implemented in MATLAB (fitglme, MathWorks). For each analysis, the dependent variable was modeled with at least fixed effects of condition (normal-hearing vs cochlear implant) and a random intercept for animal to account for repeated measurements within subjects. Model significance for fixed effects was evaluated using F-tests obtained from an ANOVA on the fitted LME model.

## Results

### iEEG recordings in normal hearing and cochlear implanted rats

We had two main goals: 1) to determine which features of auditory cortex iEEG signals were most informative about stimulus identities; and 2) to use iEEG recordings to assess cortical coding of acoustic vs electrical stimuli in normal-hearing (NH) vs bilaterally deafened + unilaterally cochlear implanted (CI) rats. Animals were first anesthetized and then a craniotomy was performed to expose the primary auditory cortex. We implanted a custom 60-channel iEEG recording array (Insanally et al., 2016; Trumpis et al., 2017) over primary auditory cortex. Seven rats were normal-hearing and presented with acoustic pure-tone stimuli (**Fig 1A**, ‘NH’). Seven rats were bilaterally deafened via cochleostomy before receiving a unilateral cochlear implant (**Fig. 1B**, ‘CI’). Peripheral sensitivity and tuning were assessed in a separate acute cohort using electrically evoked compound action potentials (ECAPs), recorded via implanted channels. Using a forward masking paradigm (Azadpour et al. 2022) we found cochlear implant stimuli were consistently tuned, cochleotopically and temporally (**Supplemental Fig. 1**).

**Figure 1.**
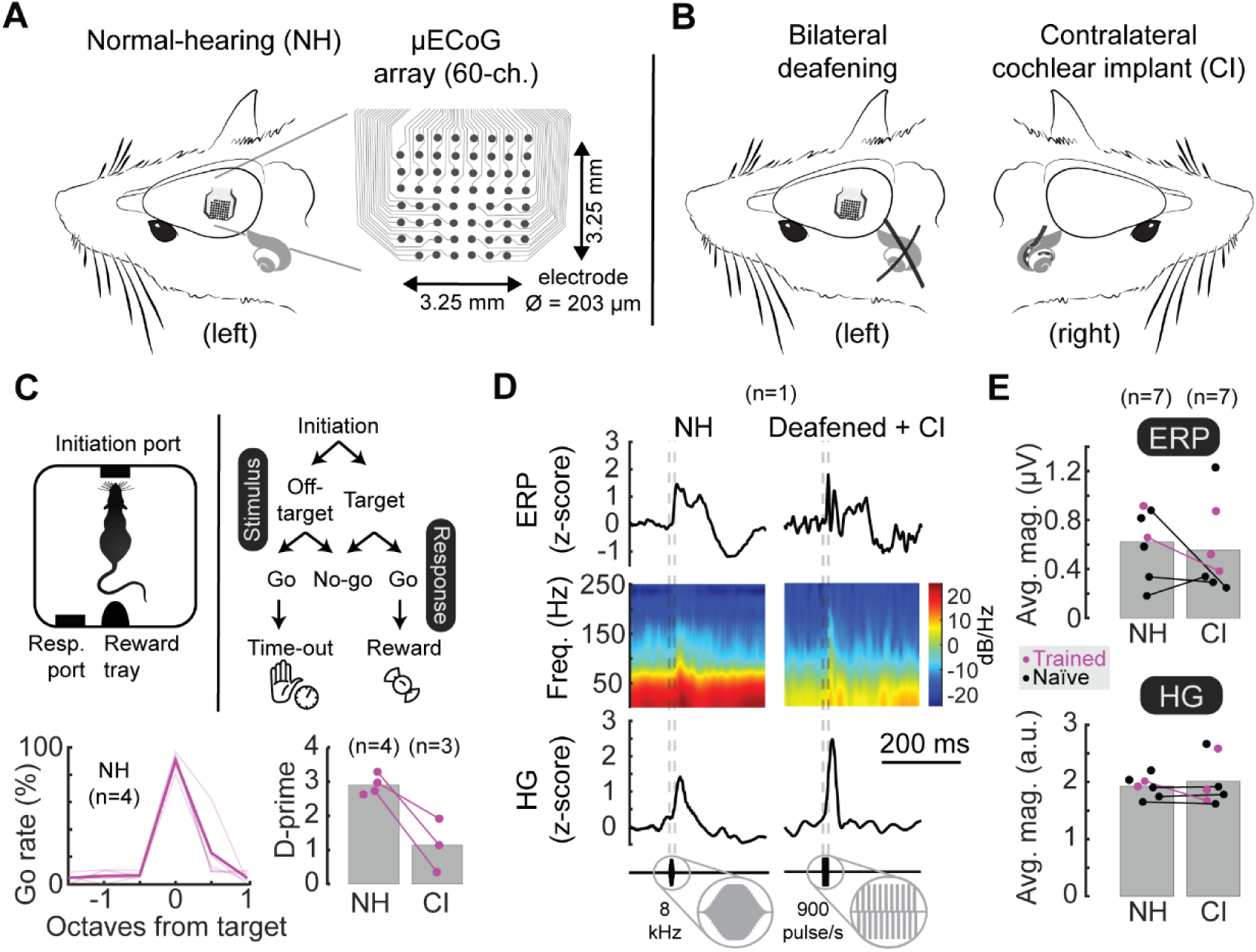
Auditory cortex iEEG responses to pure tone stimuli in normal-hearing and deafened cochlear implant rats. **A**, Schematic of 60-channel cortical surface electrode arrays, covering 3.25 x 3.25 mm on the surface of auditory cortex to record tone-evoked responses in normal-hearing (NH) trained rats. **B**, Some animals were bilaterally deafened and fitted with a unilateral cochlear implant (CI). **C**, Behavioral training of subset of animals on auditory go/no-go frequency recognition task. Animals were first trained when normal-hearing (NH, N=4, d’: 2.9±0.1) and then re-trained to respond to cochlear implant stimulation (CI, N=3, d’: 0.9±0.2). **D**, Examples of trial-averaged single-site iEEG responses (top, ERPs; middle, ERP spectrograms; bottom, HG). Clear transients were evoked by tones in normal-hearing animals (NH, left column) and in cochlear implant rats (CI, right column). **E**, iEEG response magnitude was similar between normal-hearing (NH, ERP amplitude: 1.9±0.1 µV; HG amplitude: 0.6±0.1 a.u.) and cochlear-implant rats (CI, ERP amplitude: 2.0±0.2 µV, p=0.53 compared to normal-hearing ERPs, linear mixed effects; HG amplitude: 0.6±0.1 a.u., p=0.39 compared to normal-hearing HG).

A subset of these animals from each of the two groups (N=4 normal-hearing rats, N=3 cochlear implant rats) were trained on an auditory detection and recognition go/no-go task we have used previously, both for normal hearing (Froemke et al., 2013; Carcea et al., 2017; Insanally et al., 2024) and with cochlear implants (King et al., 2016; Glennon et al., 2023). Normal-hearing rats each achieved high levels of performance, whereas the cochlear implant rats had more variable performance (**Fig. 1C**; normal-hearing behavioral performance d’: 2.9±0.1, cochlear implant performance: 0.9±0.2, one normal-hearing animal lost the implant prior to that stage of training).

The combinations of animals that underwent behavioral training and acute iEEG measurements are shown in **Supplemental Figure 2**. We acquired and then compared acute iEEG recordings of acoustic tone-evoked and electrical cochlear-implant-evoked responses in normal-hearing and cochlear implant rats. Raw responses were down-sampled and lowpass filtered to eliminate electrical artifacts from cochlear implant stimuli, and bandpass filtered again either to reveal slowly varying event-related potentials (ERPs, bandpass range: 2-100 Hz) or to reveal higher frequency high gamma oscillations (HG, bandpass range: 70-140 Hz) (**Fig. 1D**). Trial-averaged evoked responses were time-locked to the stimulus onset. Evoked response magnitudes were not measurably different between normal-hearing and cochlear implant rats (**Fig. 1E**). We found that the behavioral training (denoted purple in all figures) had no measured effect on our analyses of sensory encoding compared to responses measured in untrained rats (linear mixed effects model, p>0.3 comparing normal hearing and cochlear implant signals in untrained vs trained rats).

### A1 encoding of tones and cochlear implant electrodes are spatially organized

iEEG recordings of tone-evoked responses are spatially restricted to areas across the recording array that appeared to shift as a function of tone frequency in normal-hearing animals (**Fig. 2**). Determining the best frequency at each recording site (i.e., the stimulus evoking the maximum response overall at that location) confirmed the general tonotopic organization of auditory cortex (**Fig. 2A-E**, ERPs; **Fig. 2F-J**, HG). Tone-evoked responses were similar in expected directions and gradients (Polley et al., 2007) indicating that iEEG measurements have the spatial resolution for testing whether evoked responses are cochleotopically organized.

**Figure 2.**
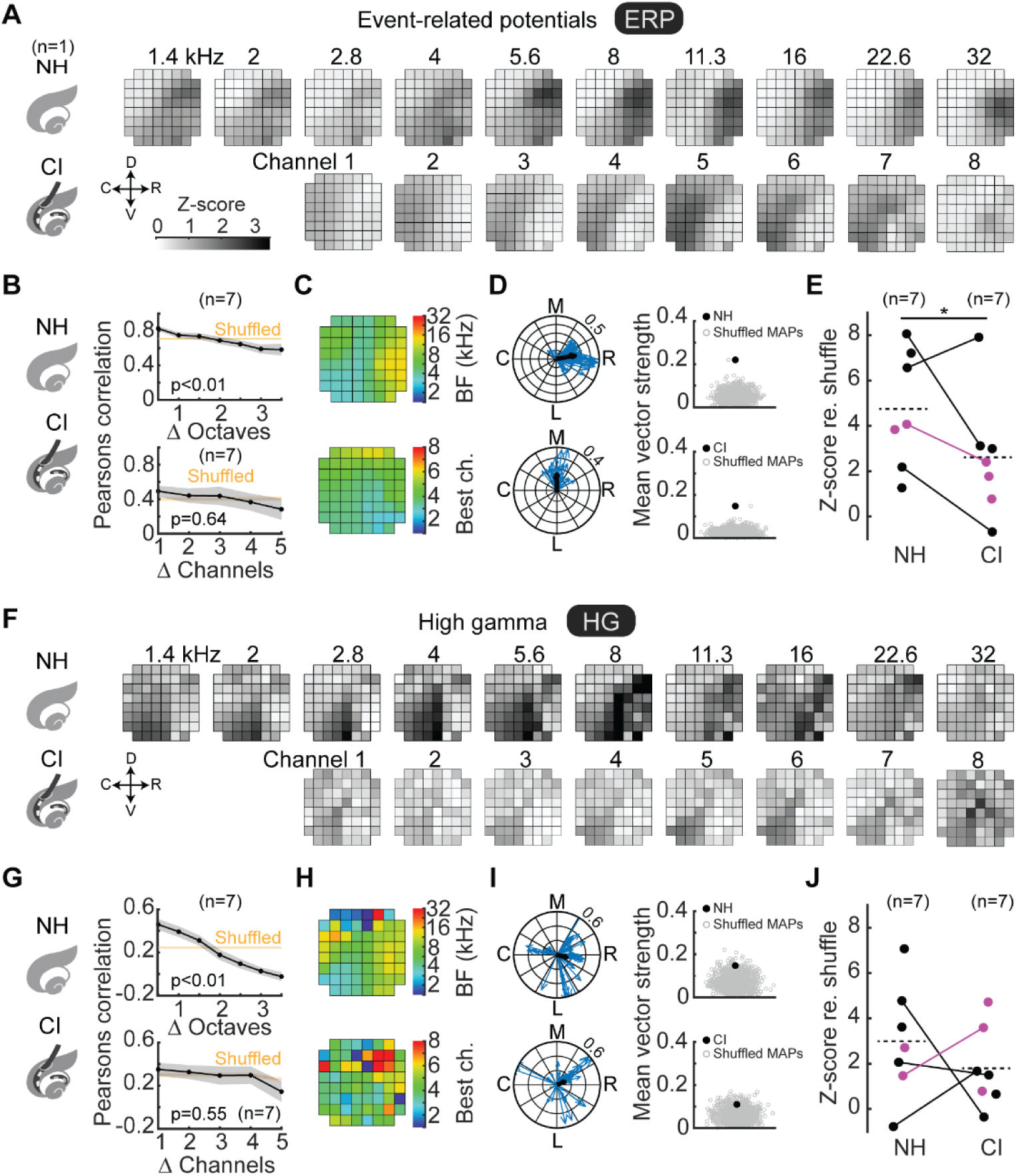
Tone-evoked and cochlear implant-evoked iEEG measurements are spatially organized. **A, F,** Trial-averaged tone-evoked event-related potentials (ERPs, panel **A**) and high gamma (HG, panel **F**) were spatially restricted to regions along the iEEG recording array; these active regions shifted as a function of stimulus. All panels represent data from the same animal (N=1), unless otherwise noted. **B, G**, Spatial correlations of evoked response areas decreased monotonically as a function of increasing stimulus separation, truncated at large separations due to low sample sizes. Evoked iEEG responses as assessed by ERPs and HG were nonrandom for normal-hearing and implanted rats (p<10^-8^ compared to all shuffled responses), but were locally tonotopic in normal-hearing rats (p<10^-4^) but not implanted rats (ERPs: p=0.5, HG: p=0.7). **C, H**, Determining the preferred stimulus of spatially-evoked activity revealed gradients shifting from high-to-low to high tone frequencies or implant channels (ERPs, panel **C**; HG, panel **H**). **D, I**, Local tonotopic gradients for each recording site plotted on a unit circle (blue) as a function of magnitude (strength of gradient) and angle (direction of gradient). The vectors across each recording site were averaged to produce an overall tonotopic vector (black) whose magnitude represents a metric of overall topography (ERPs, panel **D**; HG, panel **I**). The mean vector was plotted against the mean vector computed from n=1,000 shuffled maps (ERPs, panel **D**: normal-hearing z-score: 6.6, p<10^-10^; implanted z-score: 7.9, p<10^-14^; HG, panel **I**: normal-hearing z-score: 2.1, p=0.02; implanted z-score: 1.5, p=0.07). **E, J**, Z-scored magnitudes of non-random spatial structure across animals (ERPs, panel **E**: normal-hearing mean z-score: 4.7±1.0, implanted mean z-score: 2.6±1.0, p=0.05, linear mixed effects; HG, panel **J**: normal-hearing mean z-score: 3.0±0.9, implanted mean z-score: 2.1±0.9, p=0.48).

We then asked whether central auditory pathways would preserve some spatial ordering of cochlear implant channel identity. We found that iEEG recordings of responses evoked by individual electrodes were spatially restricted to areas across the recording array (**Fig. 2A,F**) that showed stimulus-dependent shifts in the center of cortical activation. To determine the degree of local and longer-range topography, we computed the spatial correlations as a function of stimulus separation, and compared the actual correlation coefficients to the distribution of responses when spatial locations were randomly shuffled. This analysis revealed a spatially structured but relatively coarse topographic organization (**Fig. 2B,G**; p<10^-8^ compared to all shuffled responses for ERPs and HG both for normal-hearing and implanted rats), but less sharply tonotopic relative to the tonotopic organization in normal-hearing animals assessed with pure tones (**Fig. 2B**, p<10^-4^ for ERPs and HG for normal-hearing rats; **Fig. 2G**, ERPs: p=0.64, HG: p=0.55 for implanted rats). At the extremes, the spatial correlations were always higher for small vs. large tone separations (NH 0.5 vs ≥3.5 octaves, ERP: p<10^-4^, HG: p<10^-4^ Student’s one-tailed t-test) and electrode separations (CI 1 vs ≥5 electrodes, ERP: p=0.01, HG: p=0.04).

We then reduced stimulus-evoked responses into representative stimulus maps (**Fig. 2C,H**), whose areas and orientations qualitatively match those published from similar iEEG recordings (Insanally et al. 2015), and single-unit recordings (Polley et al, 2006) albeit at coarser gradients likely due to aggregate recordings of excitatory and inhibitory activity and spatial low-pass filtering due to potentials originating far from recording sites. Direct within-animal comparisons of normal-hearing vs implant maps (N=4) indicated that cochleotopic gradients were positioned in similar orientations and directions (**Fig. 2C,H, Supplementary Fig. 3**). To quantify the degree and direction of potential spatial tonotopic or cochleotopic organization in the iEEG recordings, we computed the local gradients for each recording site (one example animal shown in **Fig. 2D**, ERPs; **Fig. 2I**, HG). Each gradient is a vector consisting of a magnitude and direction, and thus we then averaged these gradients from each recording site to get the overall cochleotopic vector, the magnitude of which is an index of spatial organization for a given animal (Romero and Hight et al. 2020). We shuffled the recording site location labels to generate putatively randomly organized maps (n=1000 random shuffles per animal) and compared the actual mean vector strength to the shuffled distributions. For this example animal, the ERP maps were significantly different from chance in terms of spatial organization (**Fig. 2D**, NH p=0.00003, CI p=0.0033), while this was more modest for the HG maps in this animal (**Fig. 2I**, NH p=0.02, CI p=0.07). This analysis revealed a similar kind of non-random spatial organization across both ERP and HG maps of normal-hearing vs cochlear implant rats (**Fig. 2E**, NH mean z-score: 4.7±1.0, CI mean z-score: 2.6±1.0, p=0.05 linear mixed effects; **Fig. 2J**, NH mean z-score: 3.0±0.9, CI mean z-score: 2.1±0.9, p=0.49), with variability across animals including some cochlear-implanted animals with effectively random global tonotopic maps. Combined with results from **Figure 2B,G**, these analyses indicate that cortical iEEG responses in normal-hearing and cochlear implant animals exhibited non-random spatial organization with coarse cochleotopic structure at the population level, but that the implant-evoked organization was substantially coarser, less sharply resolved, and more variable across animals.

### Increased variability of trial-by-trial responses evoked by cochlear implant stimulation

The results described above for **Figures 1** and **2** used trial-averaged responses to examine the spatio-temporal organization of iEEG responses. We noticed that the iEEG recordings provided satisfactory signal-to-noise ratios to measure evoked ERPs (**Fig. 3A,B**) and HG transients (**Fig. 3C,D**) on a single trial basis. Therefore, we next asked if there were differences in single-trial response variability for cochlear implant vs normal-hearing rats. We first averaged evoked responses across all recording sites to reduce measurement noise. We then computed variability of responses in the 50 ms period immediately following stimulus onset, and found that in this temporal domain, the within-animal trial-by-trial variability in evoked responses was similar for cochlear implant rats compared to normal-hearing rats irrespective of recording site (‘Temporal’; **Fig. 3B**, ERPs, linear mixed effects, p=0.054; **Fig. 3D**, HG, p=0.17).

**Figure 3.**
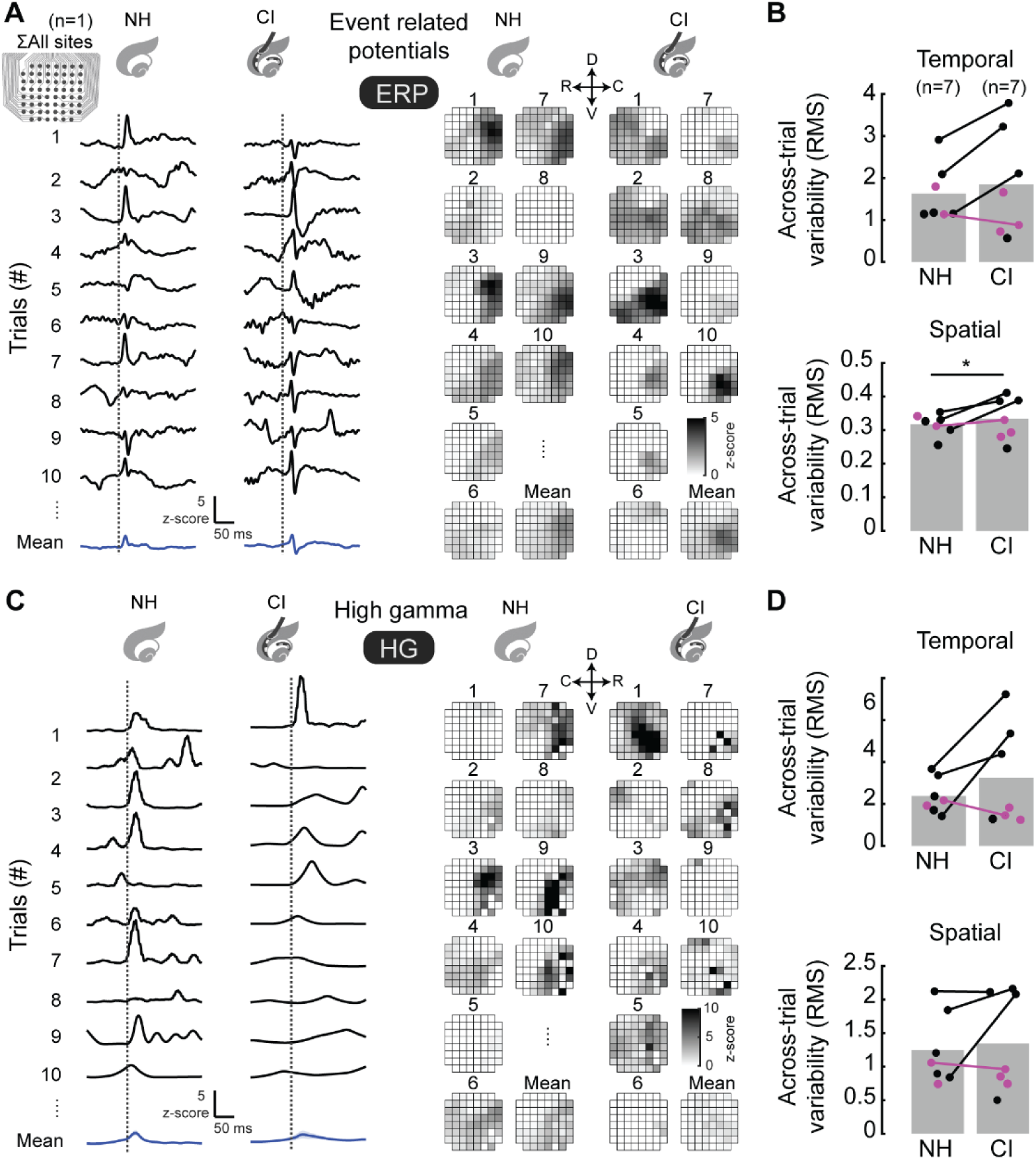
Trial-by-trial implant-evoked spatial response patterns are more variable than tone-evoked patterns. **A, C,** Trial-by-trial evoked iEEG measurements in both normal-hearing and implanted rats revealed peaks across time (left) and space (right) (ERP, panel **A**; HG, panel **C**). All panels represent data from the same animal (N=1), unless otherwise noted. **B, D,** Variability of iEEG measurements across trials (root mean square, rms) was consistently higher for spatial distributions of cochlear implant-evoked compared to tone-evoked activity, for ERPs in panel **B** (bottom, ERP spatial NH mean rms: 0.32±0.01, CI mean rms: 0.33±0.02, p=0.011, linear mixed effects) but not for temporal distributions (top, ERP temporal NH mean rms: 1.6±0.3 over all animals, CI mean rms: 1.9±0.5, for animals monitored before and after deafening, p=0.054) and HG in panel **D** (top, HG temporal NH mean rms: 2.4±0.3 over all animals, CI mean rms: 3.3±0.9, p=0.17; bottom, HG spatial NH mean rms: 1.2±0.2, CI mean rms: 1.3±0.3, p=0.35).

Then, we returned to the evoked responses for each recording site and measured the trial-by-trial evoked response peak. Next, we computed the variability of spatial distributions of evoked responses and found modestly larger within-animal trial-by-trial variability in the spatial domain of ERP responses for CI compared to normal-hearing rats (‘Spatial’; **Fig. 3B**, ERPs, linear mixed effects model, p=0.01; **Fig. 3D**, HG, p=0.35). Differences were subtle but consistent across animals (**Fig. 3B**). We conclude that similar to the trial-averaged responses in **Figure 2**, analysis of ERPs more so than HG signals could identify the increased spatial variability of evoked responses across the auditory cortex in cochlear implant animals relative to normal-hearing rats.

### Single trials encode stimulus identity in normal-hearing and cochlear implant rats

Given that trial-by-trial variability was higher for implant-evoked responses vs responses in normal-hearing rats, we wondered if this compromised decoding of cochlear implant signals in some way. We next trained a classifier to ask if we could successfully decode stimulus identity from individual trials, and examine differences of decoding accuracy for normal-hearing vs cochlear implant rats. We reduced evoked responses into the top 15 principal components via PCA (**Fig. 4A,B**). The number of components (15) was chosen as a number that modestly exceeds the number of unique stimuli identity (10 tones) while avoiding unnecessary high dimensionality. PCA scree plots (**Supplemental Fig. 4A**) showed that explained variance entered a near-linear, low-slope regime beyond this point demonstrating a similar plateau in reconstruction error (**Supplemental Fig. 4B**), indicating diminishing returns for additional components. We then trained an LDA decoder on 13 randomized trials of stimulus presentation to enable comparisons across animals that had different numbers of stimuli (Insanally et al., 2016). Computed prediction probabilities were estimated by testing the decoder on the remaining trials. Selections of the 13 training and remaining test trials were randomized (N=1,000 trials), and final prediction probabilities averaged across repeats.

**Figure 4.**
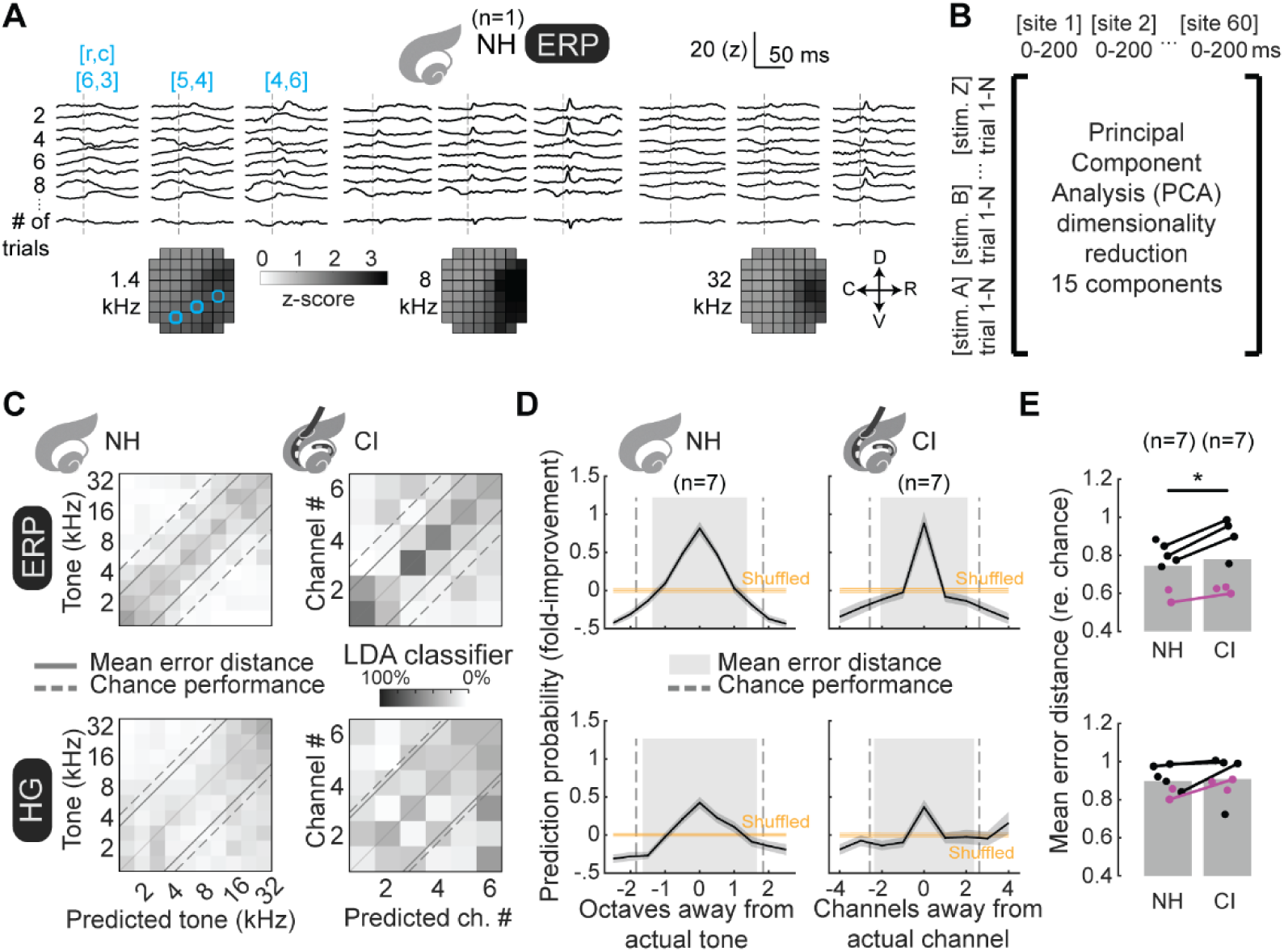
Trial-by-trial iEEG measurements encode stimulus identity. **A,** Trial-by-trial ERPs evoked by 1.4 kHz (left), 8 kHz (middle), and 32 kHz tones (right) are plotted for three recording sites (blue) and reveal discernible evoked transients and spatially restricted patterns of evoked magnitudes (stimulus onsets depicted by a dotted line). All panels represent data from the same animal (N=1), unless otherwise noted. **B,** Trial-by-trial iEEG measurements concatenated such that columns represent recording sites (spatial) and post-stimulus sampling (temporal), rows by stimulus trials. Data then reorganized by PCA and reduced to the 15 components according to magnitude of explained variance. **C,** Classification matrices are plotted as means across bootstrapped repeats (N=1,000) versions of LDA classifiers trained using 13 randomly selected trials for each stimulus, and classification predictions of single and remaining trials reveal significant prediction of stimulus identity (dashed line: chance-level error distance; solid lines: actual mean error distances). **D,** Stimulus prediction probabilities plotted across animals (black: mean, grey: s.e.m.) and as function of either octaves (normal-hearing) or channels (cochlear implant) from actual stimulus, reflecting a gradient in which adjacent stimuli share more encoded features than other stimuli. **E,** Decoder performance (mean error distance) across individual animals was somewhat worse for cochlear implant ERPs compared to tone-evoked ERPs (ERP mean error rate relative to chance for normal-hearing: 0.74±0.05, cochlear implant: 0.78±0.06; for animals assessed both first when normal-hearing and then after cochlear implantation p<10^-4^, linear mixed effects), but not for HG (HG mean error rate relative to chance for normal-hearing: 0.90±0.03, cochlear implant: 0.91±0.04; for animals assessed both first when normal-hearing and then after cochlear implantation p=0.15).

This approach successfully decoded tones and implant electrodes (**Fig. 4C**). Across rats, there were above-chance predictions of correct stimuli from single trials, with higher errors when closer to the actual stimulus (**Fig. 4D**). Computing mean error distance across trials allowed us to compare decoding performance of normal-hearing vs implanted rats, as this combines absolute prediction probability and distribution of errors while normalizing to chance-level error distances, determined by number of trained and tested stimuli. Decoders trained on responses from normal-hearing rats performed better compared to responses after implantation (**Fig. 4E**; ERP mean error distance for normal-hearing rats: 0.74±0.05, cochlear implant rats: 0.78±0.06, p<10^-4^, linear mixed effects; HG mean error distance for normal-hearing: 0.90±0.03, cochlear implant: 0.91±0.04, p=0.15).

### Single trial encoding is independently provided by both spatial and temporal cues

Which aspects of iEEG signals (spatial topography and/or temporal dynamics) contribute to stimulus decoding performance? We re-trained PCA-LDA decoders, but instead of using both spatial and temporal aspects of the full set of iEEG measurements (**Fig. 5A**), decoders were trained on the spatial-only (**Fig. 5B**) or temporal domain-only aspects (**Fig. 5C**) of evoked iEEG measurements. Decoders trained on the spatial or temporal domains only were essentially as good as predicting the stimulus as decoders trained on both aspects (**Fig. 5D,E**; mean error distance for spatial-only ERPs, normal-hearing: 0.77±0.04, implant: 0.77±0.08; for spatial-only HG, normal-hearing: 0.89±0.04, implant: 0.87±0.05; for temporal-only ERPs, normal-hearing: 0.82±0.05, implant: 0.81±0.07; for temporal-only HG, normal-hearing: 0.87±0.04, implant: 0.93±0.05; for spatial+temporal ERPs, normal-hearing: 0.74±0.04, implant: 0.86±0.06; for spatial+temporal HG, normal-hearing: 0.90±0.03, implant: 0.97±0.04). We conclude that both spatial and temporal aspects of iEEG measurements contribute to single-trial encoding of stimulus identity.

**Figure 5.**
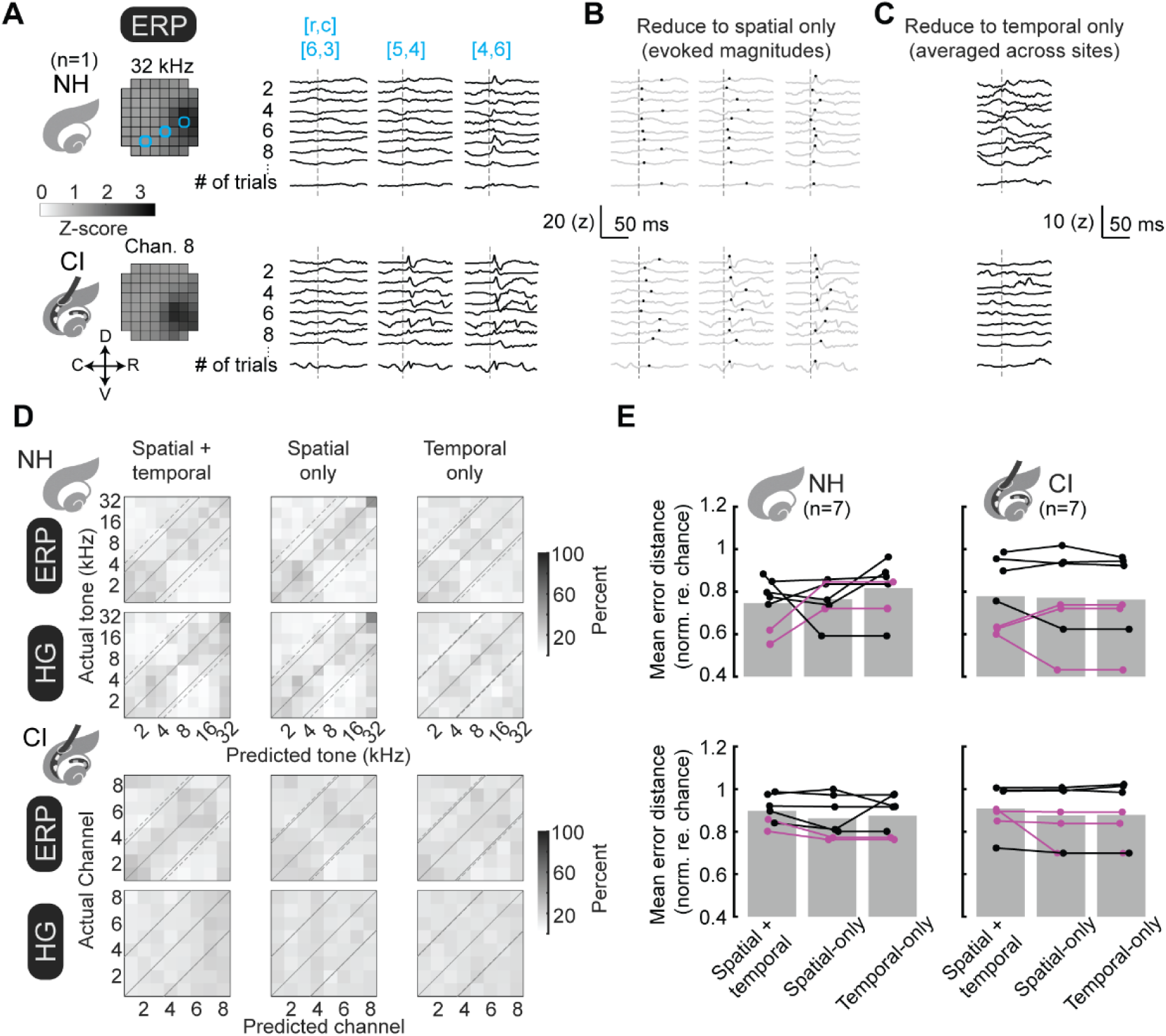
Trial-by-trial decoding from either spatial or temporal aspects of iEEG signals. **A,** Raw trial-by-trial ERPs for a single animal are plotted for 3 channels. All panels represent data from the same animal (N=1), unless otherwise noted. **B-C,** Trial-by-trial iEEG measurements are reduced into either spatial only (**B**) or temporal only (**C**) by extracting either the magnitude of evoked activity or averaging across all recording sites. **D,** Re-training PCA-LDA decoders on spatial-only (middle) or temporal-only (right) does not abolish the encoding of stimulus identity. **E,** Decoder errors (mean error distance re. chance) across animals are plotted for decoders trained on spatial+temporal, spatial-only, and temporal-only iEEG measurements. Mean error distance for spatial+temporal ERPs, normal-hearing: 0.74±0.04, implant: 0.86±0.06; for spatial+temporal HG, normal-hearing: 0.90±0.03, implant: 0.97±0.04. Mean error distance for spatial-only ERPs, normal-hearing: 0.77±0.04, implant: 0.77±0.08; for HG, normal-hearing: 0.89±0.04, implant: 0.87±0.05. Mean error distance for temporal-only ERPs, normal-hearing: 0.82±0.05, implant: 0.81±0.07; for HG, normal-hearing: 0.87±0.04, implant: 0.93±0.05.

In some cases, we found that the performance of either spatial-only or temporal-only could be somewhat better than decoders trained on both spatial and temporal aspects of iEEG measurements. This is likely due to the noise inherent in iEEG signals, such that constraining PCA reductions to one aspect of the data (either spatial or temporal) helps to eliminate variance in individual recordings.

### Reducing evoked responses using TCA preserves encoding of stimulus identity

One challenge with using PCA is that the ordinations of dimensionality reduction are unconstrained, because they are arbitrary linear combinations of the original variables. A newer approach for dimensionality reduction called tensor component analysis (TCA) explicitly enables identification of which spatial, temporal, and single-trial factors best account for signal variability, and thus drive single-trial stimulus decoding (Williams et al., 2018). Here, we used TCA instead of PCA for LDA-based decoding, constraining data reduction dimensions on the full data set to the spatial, temporal, and to the trials of iEEG measurements (**Fig. 6A**). The resulting TCA-optimizations across 15 components reveal discernable patterns of spatial and temporal factors (**Fig. 6B**). We trained and tested a LDA decoder using only the trial factors, and found that single trial predictions accurately identified the stimulus (**Fig. 6C-D**). Decoders trained on responses from normal-hearing rats had better prediction performance compared to the responses from these same animals after cochlear implantation (**Fig. 6E**; ERP mean error distance for normal-hearing rats: 0.78±0.04, for cochlear implant rats: 0.80±0.06, p<10^-5^, linear mixed-effects model; HG mean error distance for normal-hearing: 0.90±0.03, for cochlear implant: 0.88±0.05, p=0.03).

**Figure 6.**
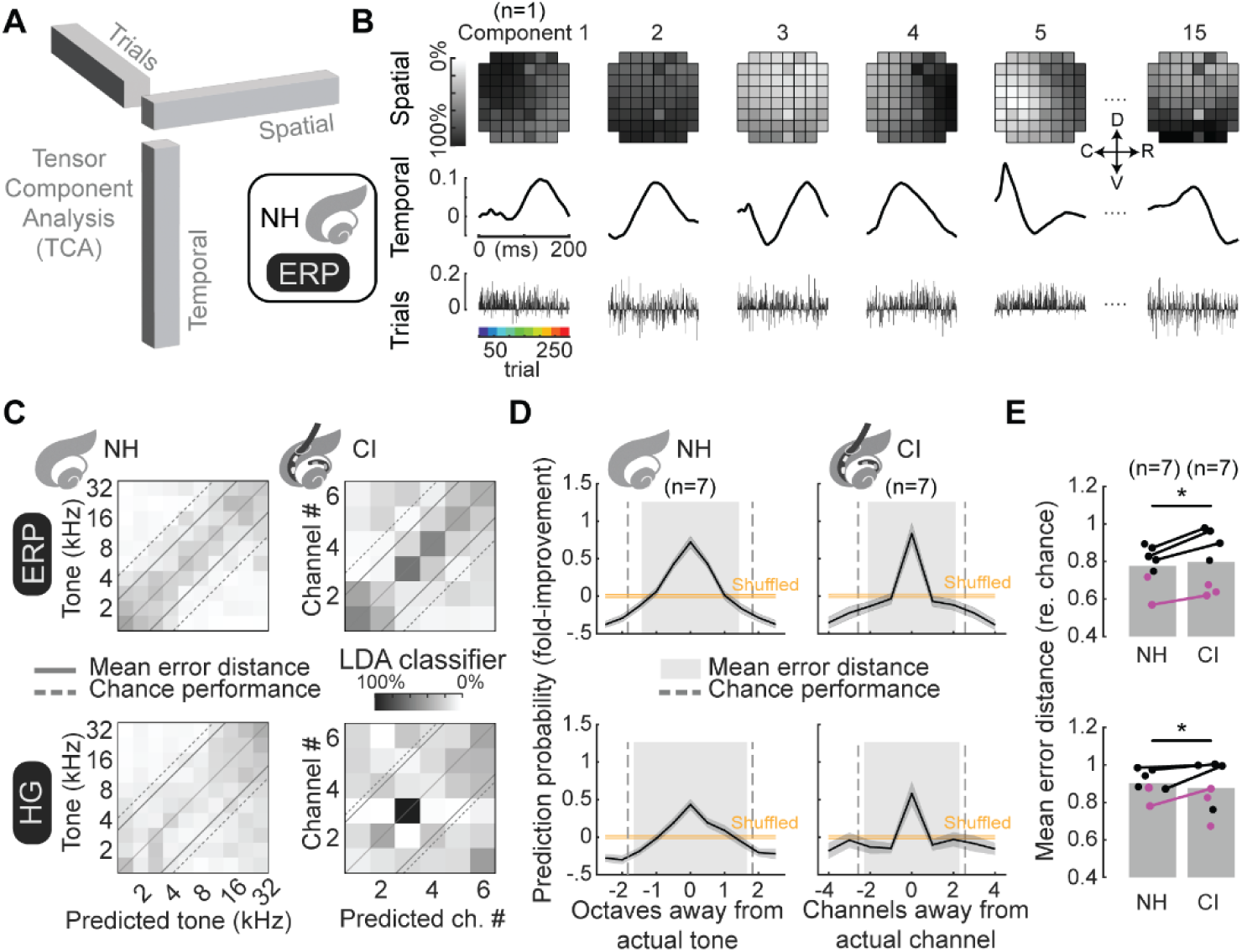
TCA-based decoding of single-trial iEEG measurements. **A,** Schematic of TCA with 3-dimensional tensors with orthogonal dimensions of spatial, temporal and trial factors. All panels represent data from the same animal (N=1), unless otherwise noted. **B,** TCA-reduced iEEG measurements of stimulus-evoked ERPs in normal-hearing rats are plotted across 15 components of spatial (top row), temporal (middle row) and trial factors (bottom row). **C,** Classification matrices showing mean decoder predictions from bootstrapped repeats (N=1,000) versions LDA classifiers trained using 13 randomly selected trials for each stimulus indicated that iEEG measurements encode stimulus identity from single stimulus presentations (dashed line: chance-level error distance; solid lines: actual mean error distances). **D,** Stimulus prediction probabilities are plotted across animals (black: mean, grey: s.e.m.) as a function of either octaves or channels away from actual stimulus, for normal-hearing and implanted rats. Correct prediction probabilities are high across all conditions and prediction errors cluster toward the actual stimuli (orange: mean and s.e.m. of decoder performance on shuffled trials; grey: mean error distance, dashed line: chance-level error distance). **E,** Decoder errors (mean error distance) across individual animals were slightly smaller for evoked iEEG measurements in normal-hearing compared to implanted rats (ERP mean error rate re. chance normal-hearing: 0.77±0.04, implanted: 0.80±0.06; HG mean error rate re. chance normal-hearing: 0.90±0.03, implanted: 0.88±0.05). Comparisons between normal-hearing and implanted iEEG measurements showed significant decrease in estimated error distances for ERP (top, linear mixed-effects model: p<10^-5^) and HG (bottom, p=0.03).

### Latent factors from unsupervised TCA revealed coarse, stimulus-related spatial structure

These analyses revealed that single-trial iEEG responses can be successfully decoded, and the TCA-based approach suggests that individual factors driving decoding might relate to specific spatiotemporal features inherent in the data. Specifically, we asked whether the TCA-reduced data contained organized spatial features, e.g., patterns of tonotopy. We next optimized TCA models of the original data (**Fig. 7A**) and then weighed the spatial factors by the strength of trial factors associated with each stimulus, resulting in spatial maps for each stimulus frequency for normal-hearing rats (**Fig. 7B**). An equivalent approach was used to generate spatial maps for each electrode in cochlear-implanted rats. The resulting reorganized spatial factors were further reduced to a preferred stimulus. Resulting TCA-maps exhibited coarse stimulus-related spatial structure (**Fig. 7C**), similar in general organization to maps extracted from ERP or HG components of the original iEEG responses. TCA-recreated spatial maps also exhibited coarse topographic organization that was non-random compared to shuffled distributions (**Fig. 7D**, p<10^-8^ compared to all shuffled responses for ERPs and HG both for normal-hearing and implanted rats). TCA models from normal-hearing HG iEEG measurements exhibited clear tonotopic organization whereas tonotopy in models from other measurements was less clear (**Fig. 7D**, normal-hearing rats ERPs: p=0.18, normal-hearing rats HG: p<10^-4^, cochlear implant rats ERPs: p=0.43, cochlear implant rats HG: p=0.40). As with analyses of **Figure 2E,J**, the mean vector strengths from TCA-derived maps indicated non-random spatial organization, and the magnitude of this organization was similar for ERP and HG measurements in normal-hearing vs cochlear implant rats (**Fig 7E**).

**Figure 7.**
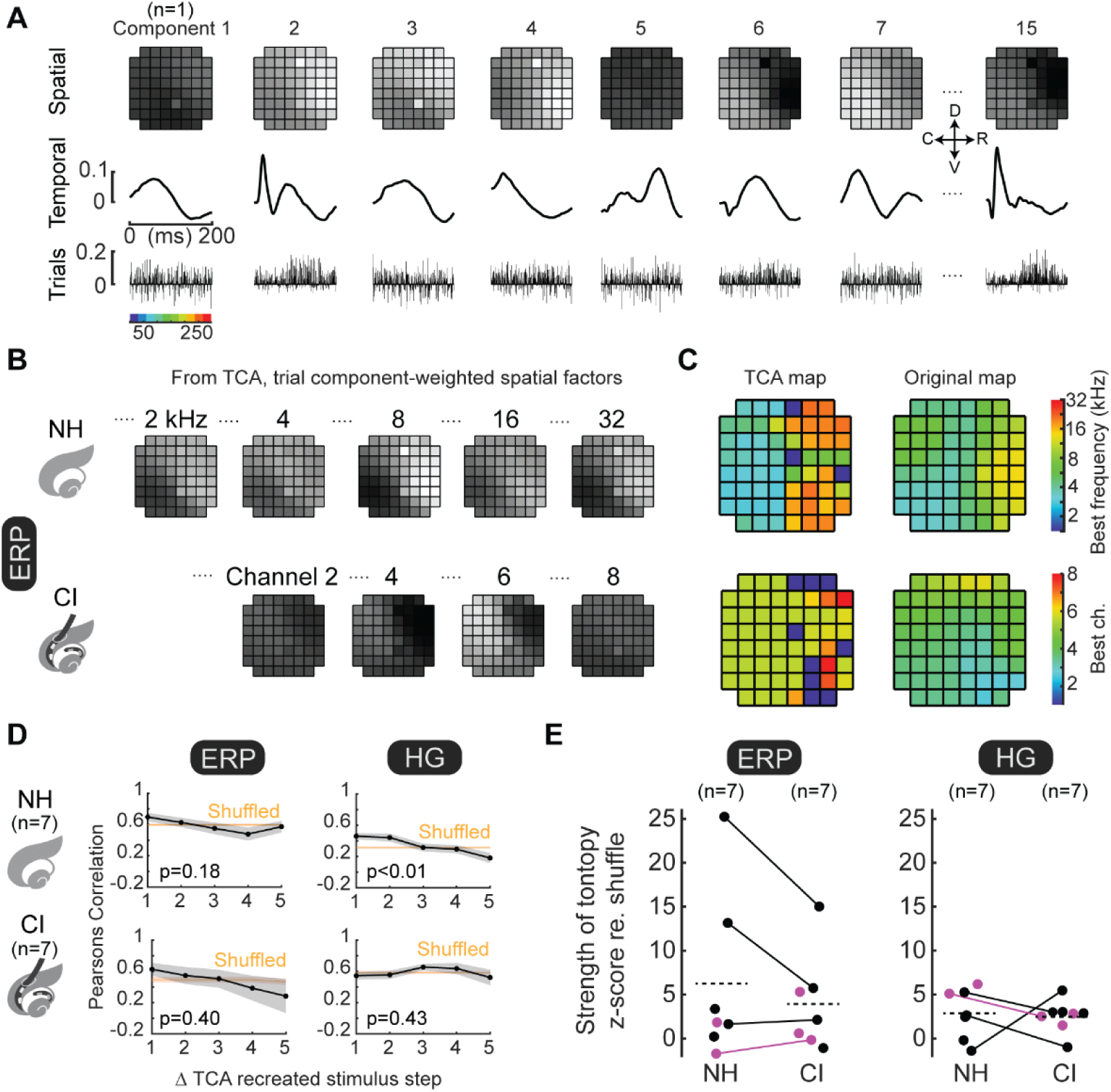
TCA data reductions reveal latent spatial factors with coarse, non-random spatial organization. **A**, TCA-based analysis of evoked neural activity, divided into 15 unique components where each component is separated into orthogonal dimensions of spatial factors (top row), temporal factors (middle row) and trial factors (bottom row). All panels represent data from the same animal (N=1), unless otherwise noted. **B**, Spatial maps for tone-evoked or electrode-evoked responses recreated from TCA reduced data. **C**, Reducing re-organized spatial factors revealed a TCA map (left) with gradients in the same location and direction as the maps reduced from raw measurements (right). **D**, Spatial correlations of evoked response areas decreased monotonically as a function of increasing stimulus separation. TCA-reduced models of evoked iEEG responses as assessed by ERPs and HG exhibited non-random spatial organization in normal-hearing and implanted rats (p<10^-8^ compared to all shuffled responses). TCA-reduced models were locally tonotopic only for HG in normal-hearing rats (normal-hearing ERPs: p=0.18, normal-hearing HG: p<10^-4^, cochlear implant ERPs: p=0.43, cochlear implant HG: p=0.40). **E**, Z-scored magnitudes of mean tonotopic vectors across animals were similar (ERPs, normal-hearing mean z-score: 6.2±3.6, cochlear implant mean z-score: 3.9±2.1, p=0.16, linear mixed effects; HG, normal-hearing mean z-score: 2.9±1.1, implanted mean z-score: 2.5±0.7, p=0.75).

### iEEG measurements from implanted rats could not be decoded from models trained on normal-hearing data

Prior analyses establish that both acoustic and implant-evoked responses are decodable on single trials, but they do not by themselves determine whether the underlying representations are shared across modalities. We next asked whether acoustic and implant-evoked responses shared a transferable spatiotemporal organization within the same animals before and after deafening and implant fitting. We used a TCA-based framework because TCA constrains the data into separable spatial, temporal, and trial factors, allowing spatial and temporal structure learned in one condition to be imposed on another. This provides a more interpretable test of cross-modal generalization than PCA, whose latent dimensions are unconstrained.

We first established a within-modality positive control by training TCA on one subset of normal-hearing trials (**Fig. 8A-E**) and then fitting withheld normal-hearing trials while holding the learned spatial and temporal factors fixed and re-optimizing only the trial factors (**Fig. 8B**). We then performed the analogous cross-modal test by fitting cochlear-implant-evoked responses using the spatial and temporal factors learned from normal-hearing data, again leaving only the trial factors free to vary (**Fig. 8F-H**). An LDA decoder trained on the normal-hearing trial factors was then tested on either the withheld normal-hearing trial factors or the cochlear-implant-derived trial factors (**Fig. 8C**). To quantify transfer without assuming any fixed mapping between predicted tones and implant electrodes, we measured how much information about actual implant-channel identity was contained in the decoder’s pattern of tone predictions. Mutual information was computed from the confusion matrix and then normalized by the maximum possible information for a matrix of that size to express information transfer as a percentage (**Fig. 8H**).

**Figure 8.**
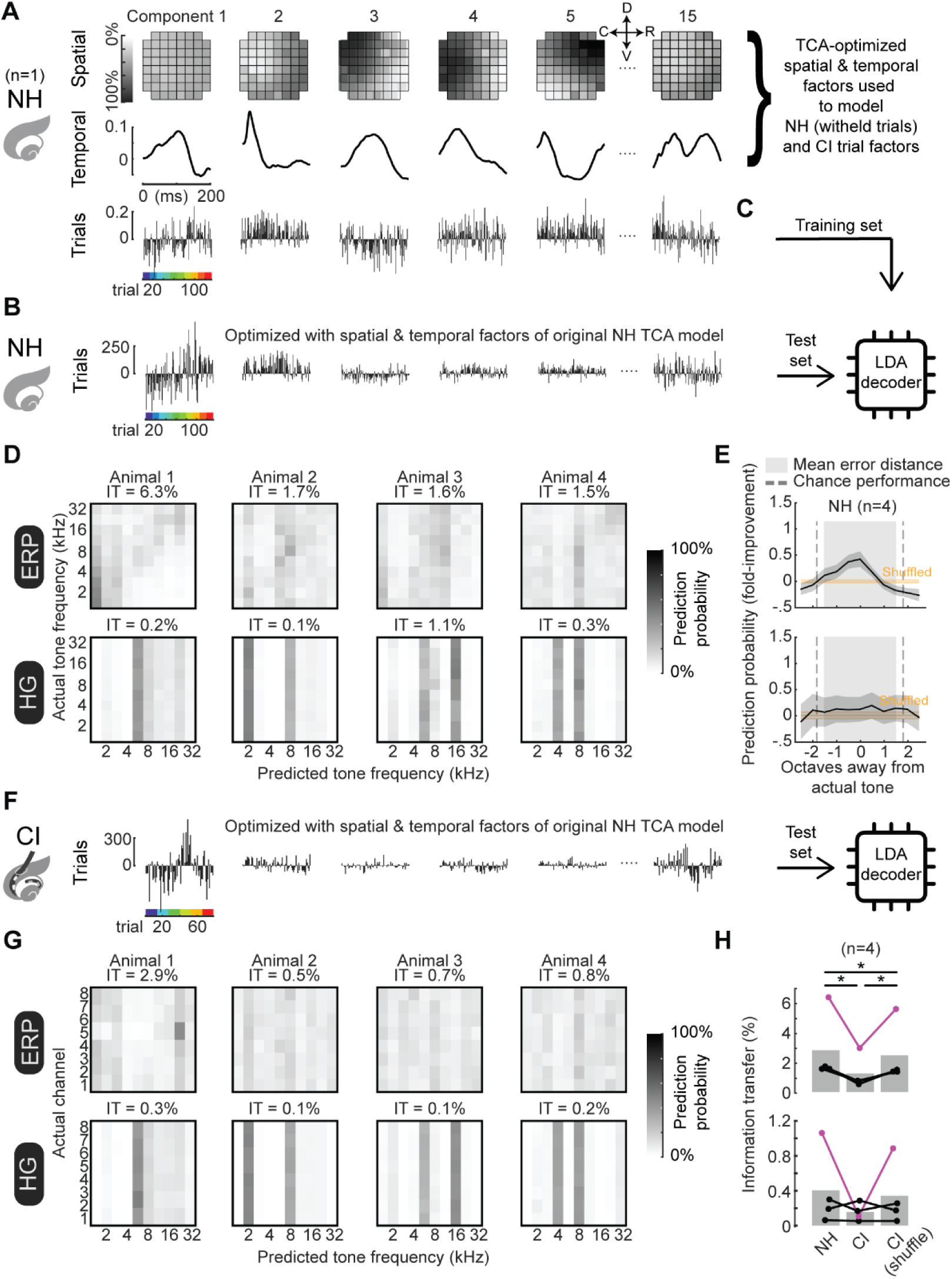
Lack of information transfer between acoustic and electrical stimulation representations in the same animals. **A,** Evoked iEEG measurements from a subset of data from normal-hearing rat reduced using TCA. All panels represent data from the same animal (N=1), unless otherwise noted. **B,** Evoked iEEG measurements from previously withheld trials from normal-hearing rat reduced using TCA, constraining the model with the spatial and temporal factors from the original TCA model, leaving only the trials as the optimizable variables. **C,** Linear discriminant analysis classifiers are trained on the trial factors extracted from the original TCA-reduced models of evoked data and then used to predict stimulus identity from the trial factors from the re-optimized TCA-reduced models of withheld normal-hearing measurements. **D,** Classification matrices are plotted as the mean decoder predictions across bootstrapped repeats (N=1,000) of linear-discriminant analysis (LDA) classifiers. Mutual information (IT) is indicated above these mean classification matrices. **E,** Stimulus prediction probabilities plotted across animals (black: mean, grey: s.e.m.) and as function of either octaves (normal-hearing) or channels (cochlear implant) from actual stimulus, reflecting a gradient in which adjacent stimuli share more encoded features than other stimuli (for ERP, but not HG). **F,** Evoked iEEG measurements in a cochlear implant rat were reduced using TCA, constraining the model with the spatial and temporal factors from the original TCA model optimized on normal-hearing data and leaving only the trials as the optimizable variables. Linear discriminant analysis classifiers were trained on the trial factors extracted from the original TCA-reduced models of evoked data in normal-hearing conditions and then used to predict stimulus identity from the trial factors from the TCA-reduced models of implant-evoked measurements. **G,** Classification matrices are plotted as the mean decoder predictions across bootstrapped repeats (N=1,000) of linear-discriminant analysis (LDA) classifiers for the normal-hearing-trained, cochlear-implant-tested condition. **H,** Mutual information (IT) is computed from each confusion matrix and divided by the maximum possible number of bits given the respective matrix dimensions. Normal-hearing-trained, implant-tested decoders reveal little-to-no information transfer (IT mean across animals, ERP: 1.2+/-0.6%; HG: 0.3+/-0.1%) compared to both the normal-hearing-only condition (IT mean across animals, ERP: 2.8+/-1.2%, p=0.04; HG: 0.4+/-0.4%, p=0.19, one-tailed t-test) and the shuffled implant-tested condition, in which rows were shuffled within columns to estimate a chance-level baseline (IT mean across animals, ERP: 2.4+/-1.0%, p=0.04; HG: 0.3+/-0.2%, p=0.21, one-tailed t-test).

We found the resulting information transfer to be low across all four animals in which we measured both tone- and implant-evoked iEEG responses (**Fig. 8H**; mean information transfer, ERPs: 1.2±0.6%; HG: 0.2±0.1%), and this was significantly lower than shuffled chance-level baseline (ERPs: 2.4±1.0%, p=0.042; HG: 0.3±0.2%, p=0.21, Student’s one-tailed t-test). Because normal-hearing-to-cochlear-implant transfer was extremely small, the shuffled baseline could equal or slightly exceed the measured value; this does not indicate recovered structure after shuffling, but rather that the observed transfer lay at or below chance. The near-zero information transfer indicates that decoders trained on tone-evoked iEEG measurements do not generalize meaningfully to implant-evoked iEEG measurements under these acute recording conditions. We repeated this analysis but with first optimizing TCA models on a subset of tone-evoked data and then constraining the spatial and temporal factors while optimizing a new TCA model of withheld tone-evoked iEEG measurements (**Fig. 8B**). Training a classifier on the original and testing the re-optimized trial factors, we found above-chance decoding accuracy and stimulus tuning (**Fig. 8C,D**). This demonstrates that this analysis approach is effective. Thus, under these acute recording conditions, acoustic-trained decoders did not generalize meaningfully to implant-evoked responses. This suggests that implant-evoked cortical activity was not simply a mildly degraded version of the acoustic representation captured by this analysis framework. Instead, the two conditions showed poor cross-modal transfer, with possible implications for perceptual similarity that remain to be tested directly.

## Discussion

Cochlear implants are the gold standard for success of brain-computer interfaces and neuroprosthetic devices to safely activate the human nervous system and effectively restore functional sensation (Hochmair et al., 2006; Wilson and Dorman, 2008; Clark, 2015; Svirsky, 2017; Glennon et al., 2020). While high levels of hearing and speech processing can be achieved by some users, outcomes remain highly variable in terms of learning rates and peak performance, especially in real-world conditions and even after controlling for age and durations of deafness (Blamey et al., 2013). Invariably, all users require an adaptation period; speech perception outcomes are initially poorer than later timepoints (Holden et al., 2013; Cusumano et al., 2017; Caswell-Midwinter et al., 2022; Stronks et al. 2025), indicating both that acute encoding of cochlear implant stimulation by the central auditory system is not optimal and that neuroplastic processes are needed for maximizing outcomes. Current and future advances in engineering and implant programming algorithms might help improve outcomes further, but there is a general belief in the field that what is now required is an understanding of how the central nervous system responds to peripheral electrical stimulation (Kral et al., 2002; Kral and Eggermont, 2007; Wilson and Dorman, 2008; Moore and Shannon, 2009).

Our findings provide the direct neural evidence that, at the time of acute cochlear implant activation, acoustic and electrical cochlear stimulation evoke cortical population responses with poor representational transfer. This addresses a longstanding question in cochlear implant research: whether electrical stimulation of the cochlea can evoke the same cortical representations as acoustic stimulation. The lack of transferable representations suggests that cochlear implant users may initially experience qualitatively different percepts that cannot be predicted from normal acoustic hearing. If so, this would challenge a foundational assumption of current cochlear implant design—that electrical stimulation should mimic the tonotopic patterns of acoustic hearing. Our data suggest that this biomimetic approach may be limited at acute activation because the cortical population responses showed poor representational transfer.

Previous studies in non-human animals have established that stimulus identity can be encoded in A1 (Bierer and Middlebrooks, 2002; Middlebrooks and Bierer, 2002; Bierer and Middlebrooks, 2004). However, many of these studies have used invasive recording methods to examine single-neuron responses to implant activation. Invasive recordings are not feasible for human cochlear implant recipients, and thus decoding methods based on purely-invasive measures may be difficult to translate into human subjects for optimizing implant programming and/or training for use of the device. While our iEEG recordings are also invasive and under anesthesia, the principles of iEEG analysis for cochlear implant responses developed here might be more applicable to non-invasive EEG recordings or similar approaches more easily performed in humans. Grid electrodes allowed us to take simultaneous advantage both of temporal and spatial features in the data, some of which were more obvious in ERPs, others were more latent in HG signals or PCA/TCA factors.

Our results show that these features could provide accurate stimulus decoding even on single trials, indicating that: 1) use of these methods in translational or clinical settings could aid personalization and optimization of brain-computer interfaces, and 2) single-trial implant responses can be adequately processed by auditory cortex and presumably downstream regions related to perception and behavior. It has been unclear to what degree the patterns of tone-evoked and implant-evoked responses in normal-hearing vs deaf subjects are similar. We found that the encoding of cochlear implant electrodes was less reliable and identifiable than tone-evoked responses in normal-hearing animals. One caveat is that direct comparisons of evoked iEEG measurements between normal-hearing and implanted rats are challenging due to fundamental differences in the stimuli (e.g., the cochleotopic separation between stimuli such as the spacing between half-octaves vs spacing between successive CI electrodes), as well as the higher degree of inter-trial variability for implant responses (which is presumably not influenced by stimulus separation). Moreover, because monopolar stimulation was delivered in a relatively smaller cochlea than in humans and recordings were made with mesoscale surface iEEG, the observed coarse cortical organization likely reflects a combination of peripheral current spread, downstream cortical pooling, and measurement resolution rather than any single limiting factor.

We also observed interesting differences in the information provided by ERPs vs HG signals. In terms of field potentials, the HG band is thought to be most directly related to neuronal spike activity (Ray and Maunsell, 2011), which may account for why HG-evoked activity was more time-locked to stimuli and more spatially tuned compared to ERPs. Some single-channel HG measurements also had higher signal-to-noise levels than ERPs. A priori, it would seem that a decoder with access to both higher signal-to-noise measurements in the temporal and spatial domains should perform better at predicting stimulus identity. Instead, we found that decoding was more accurate with ERPs than HG. One possible explanation for poorer decoding performance with HG is that HG signals are more spatially tuned, leading to a smaller number of discernable channels with clear HG activity; accordingly, we found that across iEEG grid sites, there were more significant recording sites with ERPs than HG. HG responses also have higher trial-by-trial variability and are more temporally-constrained than ERPs, and thus the ERP might be more informative in terms of decoding performance.

Finally, we found that single-trial temporal factors could suffice to provide accurate decoding of implant channels. This suggests that preservation of clear cochleotopy in deaf subjects may not be a pre-requisite for successful implant use. Rather, it is likely that reductions in trial-by-trial and inter-neuronal variability might be more important as a mechanism for improving outcomes. This means that the neural responses to neuroprosthetic stimulation and use of brain-computer interfaces need not fully recapitulate responses to other modalities (e.g., acoustic stimulation) in space and time. These findings are consistent with the possibility that cochlear implant stimulation engages an alternative representation in auditory cortex, rather than a simple degraded version of acoustic encoding. A key limitation, however, is that these recordings were obtained acutely, under anesthesia, and before prolonged experience with the implant. We therefore cannot determine whether the poor overlap observed here is transient, whether cortical representations become more aligned with chronic use, whether downstream readout adapts to a novel code, or whether both processes occur. Longitudinal recordings during chronic implant use will be required to distinguish between these possibilities. Previously we showed that deafened rats could learn to use a cochlear implant to perform an auditory task, and that changes in cortical excitatory and inhibitory synapses related to implant use outcomes. Our results suggest that refinement particularly of inhibitory cortical responses might help increase inter-neuronal variance at the population level to increase spatial organization, while decreasing intra-trial variability at the single neuron level to enhance temporal organization and improve auditory perception.

## Acknowledgements

We thank Badr Albanna, Elad Sagi, and Sam Zheng for comments, discussions, and technical assistance. This work was funded by the National Institute on Deafness and Other Communication Disorders (K99DC021727 to A.E.H., F30DC015170 to J.K.S., and R01DC012557 to R.C.F. and M.A.S.), the National Center for Advancing Translational Sciences (TL1TR001447 to A.E.H.), and the Charles H. Revson Foundation (to A.E.H.); the statements made and views expressed, however, are solely the responsibility of its authors. The current affiliation for Julia K. Scarpa is Weill Cornell Medicine, New York, NY 10065; for Yew-Song Cheng is Massachusetts Eye and Ear Infirmary, Harvard Medical School, Boston MA, 02114; for Michael Trumpis is Paradromics Inc., Austin TX 78759.

## Figures

**Supplemental Figure 1.**
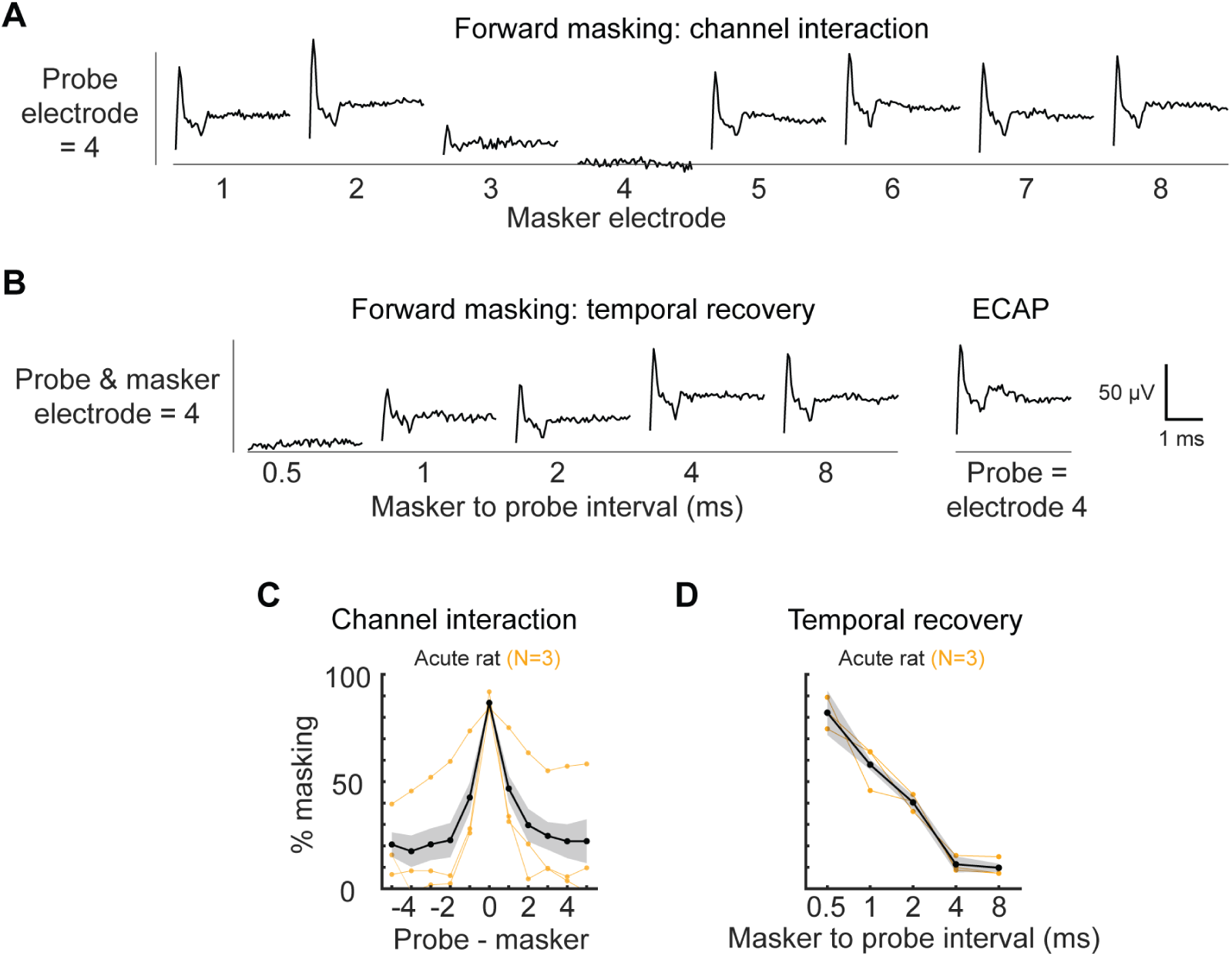
ECAP measurements of spatio-temporal cochlear implant tuning. **A,** Forward masking was performed to assess the spatial tuning of cochlear stimuli in the cochlea via ECAPs measured by adjacent cochlear implant electrodes. These measurements were made in a separate group of animals (N=3) than those used for iEEG recordings. **B,** Forward masking was performed to assess temporal tuning of cochlear implant stimuli in the cochlea via ECAPs measured by adjacent cochlear implant electrodes. The masker stimulus was delivered by the same electrode as the probe. **C,** Spatial tuning functions were averaged across all probe electrodes in N=3 animals (black, mean; gray: s.e.m.; orange, average of individual subjects). **D,** Temporal tuning functions were averaged across all probe electrodes and N=3 animals (black, mean; gray, s.e.m.; orange, average of individual subjects).

**Supplemental Figure 2.**
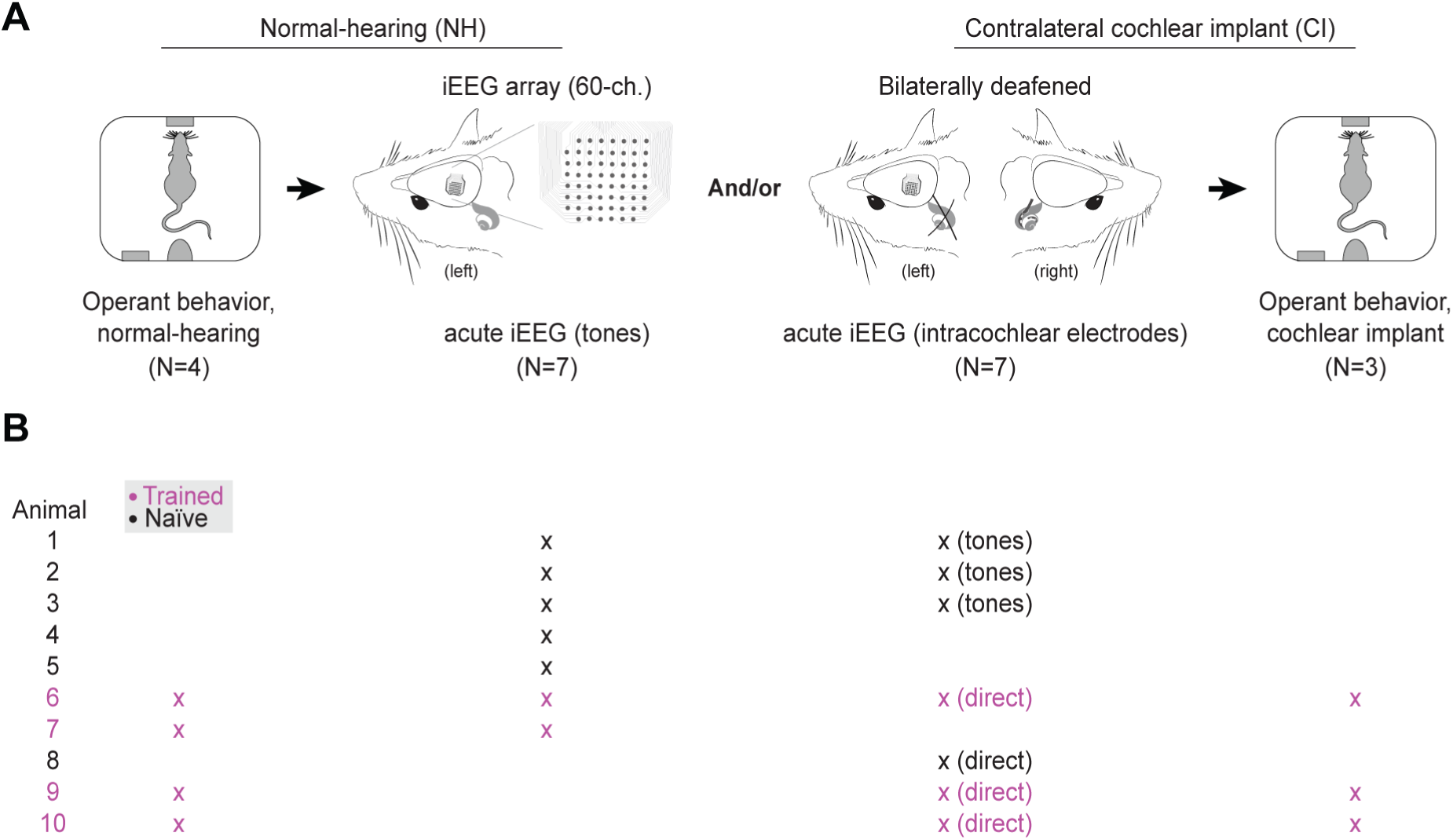
Experimental timeline and animal tracker. **A,** Rats underwent a combination of behavioral training (when typically-hearing), iEEG recordings, and then behavioral testing (after deafening) in a specific sequence, but not all animals included here participated in every phase. **B,** iEEG measurements were obtained in a total of N=10 animals. In a subset of these animals (N=4), we also obtained iEEG measurements in response to both acoustic tones before deafening and cochlear implant stimulation after deafening. In addition to iEEG recordings, we also behaviorally trained either prior (tones) or after iEEG recordings with cochlear implant stimuli.

**Supplemental Figure 3.**
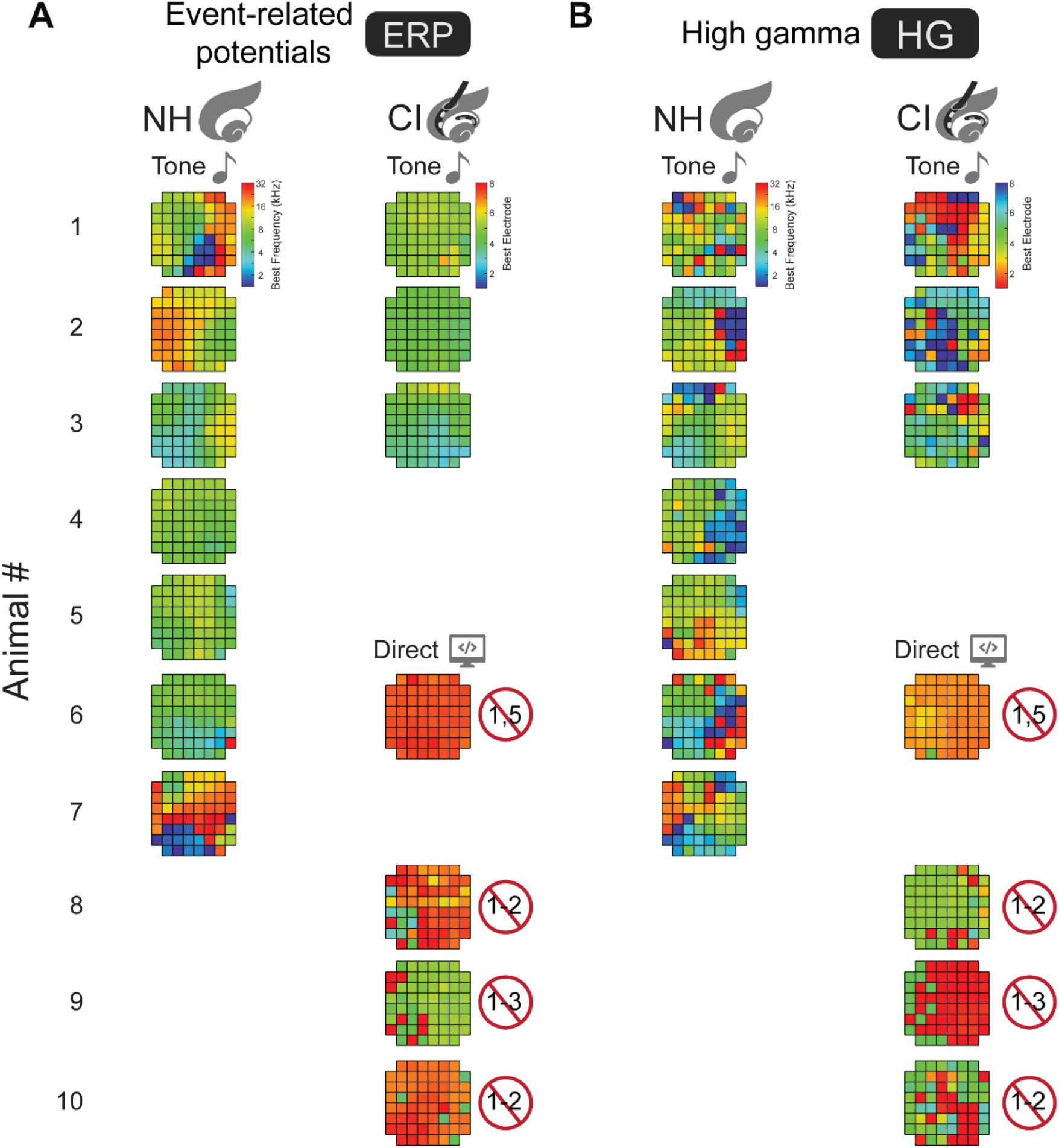
Maps of stimulus preferences for tone- or cochlear implant-evoked iEEG measurements. **A,** Maps of preferred stimuli are derived from event-related potentials (ERP) for 10 animals, including tone evoked iEEG measurements (left column) and cochlear implant-evoked iEEG\ measurements (right column). For the cochlear implant, single channel stimuli were either delivered via tones presented to the cochlear implant processor (top 3 maps) and or single channel stimuli were triggered by a direct connection between computer and the cochlear implant processor (bottom 3 maps). In some cochlear-implanted animals, apical electrodes were non-functional. The numbers of these non-functional electrodes are denoted with a superimposed red circle and backslash.

**Supplemental Figure 4:**
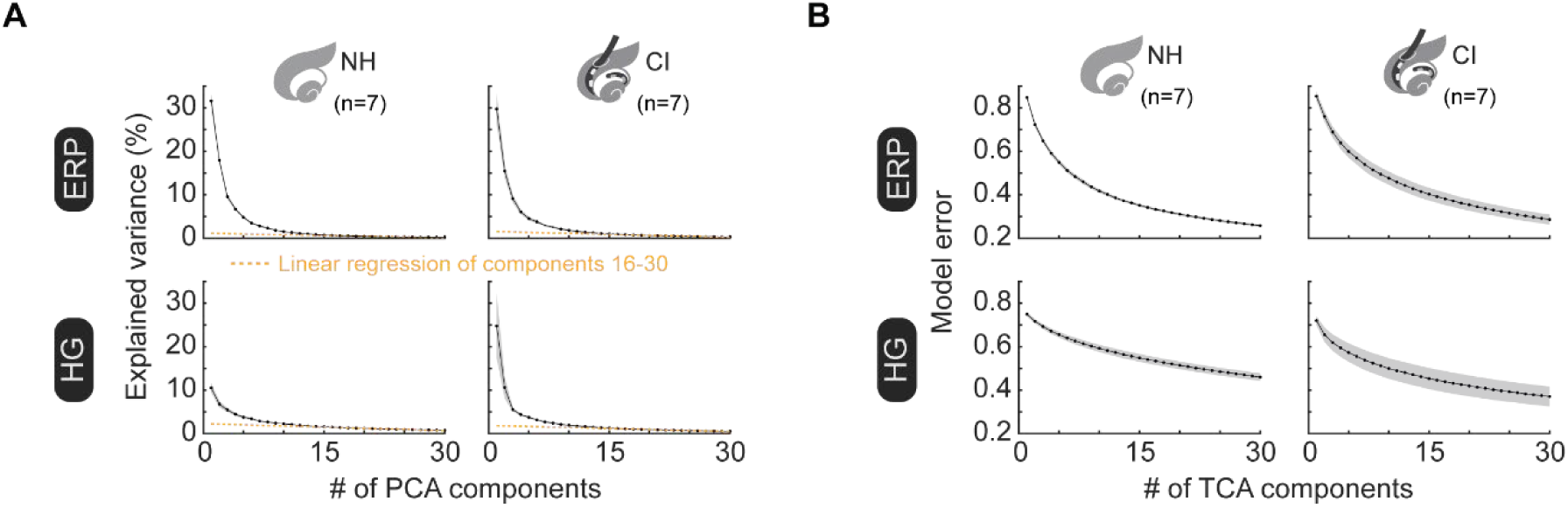
Numbers of components for PCA and TCA models. **A.** The means (+/- s.e.m.) of explained variance per component for PCA models across all ERP data (top row) decrease rapidly from 30 percent to near zero within 30 components. The means (+/- s.e.m.) of explained variance per component for PCA models for HG data (bottom row) and decrease from 12 percent rapidly to a steady decline within 30 components. Orange dashed line: linear regression of components 16-30 are plotted. **B.** The means (+/- s.e.m.) of model error for TCA models of ERP (top row) and HG (bottom row) decrease as a function of number of components.

## References

Abdi-Sargezeh B, Oswal A, Sanei S (2023) Mapping scalp to intracranial EEG using generative adversarial networks for automatically detecting interictal epileptiform discharges. In: 2023 IEEE Statistical Signal Processing Workshop (SSP), 710–714.

Adenis, V, et al. “Asymmetric Pulses Delivered by a Cochlear Implant Allow a Reduction in Evoked Firing Rate and in Spatial Activation in the Guinea Pig Auditory Cortex.” Hearing Research [Netherlands], vol. 447, no. 109027, June 2024, 10.1016/j.heares.2024.109027.

Ayton LN, Barnes N, Dagnelie G, Fujikado T, Goetz G, Hornig R, Jones BW, Muqit MMK, Rathbun DL, Stingl K, Weiland JD, Petoe MA (2020) An update on retinal prostheses. Clin Neurophysiol 131:1383–1398.

Azadpour M, McKay CM (2012) A psychophysical method for measuring spatial resolution in cochlear implants. J Assoc Res Otolaryngol 13:145–157.

Bader BW, Kolda TG, Others a (2023) Tensor Toolbox for MATLAB. In, 3.6 Edition.

Beynon AJ, Luijten BM, Mylanus EAM (2021) Intracorporeal cortical telemetry as a step to automatic closed-loop EEG-based CI fitting: A proof of concept. Audiol Res 11:691–705.

Bierer JA, Middlebrooks JC (2002) Auditory cortical images of cochlear-implant stimuli: dependence on electrode configuration. J Neurophysiol 87:478–492.

Bierer JA, Middlebrooks JC (2004) Cortical responses to cochlear implant stimulation: channel interactions. J Assoc Res Otolaryngol 5:32–48.

Blamey P et al. (2013) Factors affecting auditory performance of postlinguistically deaf adults using cochlear implants: an update with 2251 patients. Audiol Neurootol 18:36–47.

Carcea I, Insanally MN, Froemke RC (2017) Dynamics of auditory cortical activity during behavioural engagement and auditory perception. Nat Commun 8:14412.

Caswell-Midwinter B, Doney EM, Arjmandi MK, Jahn KN, Herrmann BS, Arenberg JG (2022) The relationship between impedance, programming and word recognition in a large clinical dataset of cochlear implant recipients. Trends Hear 26:23312165211060983.

Chang EF (2015) Towards large-scale, human-based, mesoscopic neurotechnologies. Neuron 86:68–78.

Clark GM (2006) The multiple-channel cochlear implant: the interface between sound and the central nervous system for hearing, speech, and language in deaf people-a personal perspective. Philos Trans R Soc Lond B Biol Sci 361:791–810.

Clark GM (2015) The multi-channel cochlear implant: Multi-disciplinary development of electrical stimulation of the cochlea and the resulting clinical benefit. Hear Res 322: 4–13.

Cusumano C, Friedmann DR, Fang Y, Wang B, Roland JT, Jr., Waltzman SB (2017) Performance plateau in prelingually and postlingually deafened adult cochlear implant recipients. Otol Neurotol 38:334–338.

Dawson J, Pierce D, Dixit A, Kimberley TJ, Robertson M, Tarver B, Hilmi O, McLean J, Forbes K, Kilgard MP, Rennaker RL, Cramer SC, Walters M, Engineer N (2016) Safety, feasibility, and efficacy of vagus nerve stimulation paired with upper-limb rehabilitation after ischemic stroke. Stroke 47:143–150.

Djourno A, Eyries C (1957) [Auditory prosthesis by means of a distant electrical stimulation of the sensory nerve with the use of an indwelt coiling]. Presse Med (1893) 65:1417.

Eddington DK, Dobelle WH, Brackmann DE, Mladejovsky MG, Parkin JL (1978) Auditory prostheses research with multiple channel intracochlear stimulation in man. Ann Otol Rhinol Laryngol 87:1–39.

Fallon JB, Shepherd RK, Irvine DRF (2014) Effects of chronic cochlear electrical stimulation after an extended period of profound deafness on primary auditory cortex organization in cats. Eur J Neurosci 39:811–820.

Froemke RC, Carcea I, Barker AJ, Yuan K, Seybold BA, Martins ARO, Zaika N, Bernstein H, Wachs M, Levis PA, Polley DB, Merzenich MM, Schreiner CE (2013) Long-term modification of cortical synapses improves sensory perception. Nat Neurosci 16:79–88.

Fukushima M, Chao ZC, Fujii N (2015) Studying brain functions with mesoscopic measurements: Advances in electrocorticography for non-human primates. Curr Opin Neurobiol 32:124–131.

Glennon E, Svirsky MA, Froemke RC (2020) Auditory cortical plasticity in cochlear implant users. Curr Opin Neurobiol 60:108–114.

Glennon E, Valtcheva S, Zhu A, Wadghiri YZ, Svirsky MA, Froemke RC (2023) Locus coeruleus activity improves cochlear implant performance. Nature 613:317–323.

Glennon E, Carcea I, Martins ARO, Multani J, Shehu I, Svirsky MA, Froemke RC (2019) Locus coeruleus activation accelerates perceptual learning. Brain Res 1709:39–49.

Hays SA, Rennaker RL, Kilgard MP (2013) Targeting plasticity with vagus nerve stimulation to treat neurological disease. Prog Brain Res 207:275–299.

Hochmair ES, Hochmair-Desoyer IJ (1981) An implanted auditory eight channel stimulator for the deaf. Med Biol Eng Comput 19:141–148.

Hochmair I, et al. (2006) MED-EL cochlear implants: State of the art and a glimpse into the future. Trends Amplif 10:201–219.

Holden LK, Finley CC, Firszt JB, Holden TA, Brenner C, Potts LG, Gotter BD, Vanderhoof SS, Mispagel K, Heydebrand G, Skinner MW (2013) Factors affecting open-set word recognition in adults with cochlear implants. Ear Hear 34:342–360.

House WF, Urban J (1973) Long term results of electrode implantation and electronic stimulation of the cochlea in man. Ann Otol Rhinol Laryngol 82:504–517.

Insanally M, Trumpis M, Wang C, Chiang C-H, Woods V, Palopoli-Trojani K, Bossi S, Froemke RC, Viventi J (2016) A low-cost, multiplexedμECoG system for high-density recordings in freely moving rodents. J Neural Eng 13:026030–26030.

Insanally MN, Albanna BF, Toth J, DePasquale B, Fadaei SS, Gupta T, Lombardi O, Kuchibhotla K, Rajan K, Froemke RC (2024) Contributions of cortical neuron firing patterns, synaptic connectivity, and plasticity to task performance. Nat Commun 15:6023.

Janiukstyte V, Owen TW, Chaudhary UJ, Diehl B, Lemieux L, Duncan JS, de Tisi J, Wang Y, Taylor PN (2023) Normative brain mapping using scalp EEG and potential clinical application. Sci Rep 13:13442.

Jasper H, Penfield W (1949) Electrocorticograms in man: Effect of voluntary movement upon the electrical activity of the precentral gyrus. Archiv für Psychiatrie und Nervenkrankheiten 183:163–174.

Johnson KA et al. (2024) Proceedings of the 11th Annual Deep Brain Stimulation Think Tank: pushing the forefront of neuromodulation with functional network mapping, biomarkers for adaptive DBS, bioethical dilemmas, AI-guided neuromodulation, and translational advancements. Front Hum Neurosci 18:1320806.

Johnson LA, Della Santina CC, Wang X (2016) Selective neuronal activation by cochlear implant stimulation in auditory cortex of awake primate. J Neurosci 36:12468–12484.

Keating P, Nodal FR, Gananandan K, Schulz AL, King AJ (2013) Behavioral sensitivity to broadband binaural localization cues in the ferret. J Assoc Res Otolaryngol 14:561–572.

King J, Shehu I, Roland JT, Jr., Svirsky MA, Froemke RC (2016) A physiological and behavioral system for hearing restoration with cochlear implants. J Neurophysiol 116:844–858.

Klinke R, Kral A, Heid S, Tillein J, Hartmann R (1999) Recruitment of the auditory cortex in congenitally deaf cats by long-term cochlear electrostimulation. Science 285:1729–1733.

Kolda TG, Bader BW (2009) Tensor Decompositions and Applications. SIAM Review 51:455–500.

Kral A, et al. (2002) Hearing after congenital deafness: Central auditory plasticity and sensory deprivation. Cereb Cortex 12:797–807.

Kral A, Eggermont JJ (2007) What’s to lose and what’s to learn: Development under auditory deprivation, cochlear implants and limits of cortical plasticity. Brain Res Rev 56:259–269.

Martins ARO, Froemke RC (2015) Coordinated forms of noradrenergic plasticity in the locus coeruleus and primary auditory cortex. Nat Neurosci 18:1483–1492.

McDermott HJ (2004) Music perception with cochlear implants: a review. Trends Amplif 8:49–82.

Merzenich MM, Knight PL, Roth GL (1973) Cochleotopic organization of primary auditory cortex in the cat. Brain Res 63:343–346.

Mesgarani N, Cheung C, Johnson K, Chang EF (2014) Phonetic feature encoding in human superior temporal gyrus. Science 343:1006–1010.

Michelson RP (1971) The results of electrical stimulation of the cochlea in human sensory deafness. Ann Otol Rhinol Laryngol 80:914–919.

Middlebrooks JC, Bierer JA (2002) Auditory cortical images of cochlear-implant stimuli: coding of stimulus channel and current level. J Neurophysiol 87:493–507.

Moore DR, RV Shannon (2009) Beyond cochlear implants: awakening the deafened brain. Nat Neurosci 12:686–691.

Norman-Haignere SV, Long LK, Mesgarani N, Devinsky O, Flinker A, Doyle W, Irobunda I, Merricks EM, Schevon CA, Feldstein NA, McKhann GM (2022) Multiscale temporal integration organizes hierarchical computation in human auditory cortex. Nat Hum Behav 6:455–469-469.

Nourski KV, Etler CP, Brugge JF, Oya H, Kawasaki H, Reale RA, Abbas PJ, Brown CJ, Howard MA (2013) Direct recordings from the auditory cortex in a cochlear implant user. J Assoc Res Otolaryngol 14:435–450.

Polley DB, Read HL, Storace DA, Merzenich MM (2007) Multiparametric auditory receptive field organization across five cortical fields in the albino rat. J Neurophysiol 97:3621–3638.

Ray S, Maunsell JH (2011) Different origins of gamma rhythm and high-gamma activity in macaque visual cortex. PLoS Biol 9:e1000610.

Simmons FB, Epley JM, Lummis RC, Guttman N, Frishkopf LS, Harmon LD, Zwicker E (1965) Auditory Nerve: Electrical Stimulation in Man. Science 148:104–106.

Stronks, HC, Arendsen TS, Veenstra M, Boermans P-PBM, Briaire JJ, Frijns JHM (2025) Effects of preoperative factors on the learning curves of postlingual cochlear implant recipients. Ear Hear (in press).

Svirsky M (2017) Cochlear implants and electronic hearing. Physics Today 70:52–58.

Tang C, Hamilton LS, Chang EF (2017) Intonational speech prosody encoding in the human auditory cortex. Science 357:797–801.

Trumpis M, Insanally M, Zou J, Elsharif A, Ghomashchi A, Sertac Artan N, Froemke RC, Viventi J (2017) A low-cost, scalable, current-sensing digital headstage for high channel count muECoG. J Neural Eng 14:026009.

Vermeire K, Landsberger DM, Schleich P, Van de Heyning PH (2013) Multidimensional scaling between acoustic and electric stimuli in cochlear implant users with contralateral hearing. Hear Res 306:29–36.

Vollmer M, Beitel RE (2011) Behavioral training restores temporal processing in auditory cortex of long-deaf cats. J Neurophysiol 106:2423–2436.

Walzl EM, Woolsey CN (1946) Effects of cochlear lesions on click responses in the auditory cortex of the cat. Bull Johns Hopkins Hosp 79:309–319.

Williams AH, Kim TH, Wang F, Vyas S, Ryu SI, Shenoy KV, Schnitzer M, Kolda TG, Ganguli S (2018) Unsupervised Discovery of Demixed, Low-Dimensional Neural Dynamics across Multiple Timescales through Tensor Component Analysis. Neuron 98:1099–1115.

Wilson BS, Finley CC, Lawson DT, Wolford RD, Eddington DK, Rabinowitz WM (1991) Better speech recognition with cochlear implants. Nature 352:236–238.

Wilson BS, MF Dorman (2008) Cochlear implants: A remarkable past and a brilliant future. Hear Res 242:3–21.

Woods V, Trumpis M, Bent B, Palopoli-Trojani K, Chiang C-H, Wang C, Yu C, Insanally MN, Froemke RC, Viventi J (2018) Long-term recording reliability of liquid crystal polymer µECoG arrays. J Neural Eng 15:066024.

Zeng FG (2022) Celebrating the one millionth cochlear implant. JASA Express Lett 2:077201.

